# Separase cleaves the kinetochore protein Meikin to direct the meiosis I/II transition

**DOI:** 10.1101/2020.11.29.402537

**Authors:** Nolan K Maier, Jun Ma, Michael A Lampson, Iain M Cheeseman

## Abstract

To generate haploid gametes, germ cells undergo two consecutive meiotic divisions requiring key changes to the cell division machinery. Here, we explore the regulatory mechanisms that differentially control meiotic events. We demonstrate that the protease Separase rewires key cell division processes at the meiosis I/II transition by cleaving the meiosis-specific protein Meikin. In contrast to cohesin, which is inactivated by Separase proteolysis, cleaved Meikin remains functional, but results in a distinct activity state. Full-length Meikin and the C-terminal Meikin Separase-cleavage product both localize to kinetochores, bind to Plk1 kinase, and promote Rec8 cleavage, but our results reveal distinct roles for these proteins in controlling meiosis. Mutations that prevent Meikin cleavage or that conditionally inactivate Meikin at anaphase I both result in defective meiosis II chromosome alignment. Thus, Separase cleavage of Meikin creates an irreversible molecular switch to rewire the cell division machinery at the meiosis I/II transition.

## Introduction

The generation of haploid germ cells from diploid precursors requires a specialized cell division, termed meiosis, in which two successive rounds of chromosome segregation occur without undergoing intervening DNA replication (Miller et al., 2013). In most organisms, the first round of chromosome segregation during meiosis I reduces the chromosome number by half, requiring three major modifications relative to the mitotic cell division program. First, homologous chromosomes pair and become physically linked through synapsis and recombination. Second, instead of binding to opposing spindle poles, the kinetochores of sister chromatids co-orient to connect to microtubules from the same spindle pole. Third, cohesin complexes adjacent to centromeres are retained through to the second meiotic division even as cohesin along the chromosome arms is removed, allowing sister chromatids to remain associated and co-segregate to the same pole. Each of these modifications to meiosis I requires the activity of meiosis-specific proteins whose expression is restricted to germ cells. For example, the vertebrate factor Meikin (meiosis-specific kinetochore protein) plays important roles in meiosis I sister kinetochore co-orientation and the protection of centromeric cohesion (Kim et al., 2015a), with *Meikin*-null mice exhibiting meiotic defects and sterility. Current functional evidence suggests that Meikin plays an analogous role to that of Spo13 and Moa1 in yeast meiosis (Galander et al., 2019b; Kim et al., 2015a; Miyazaki et al., 2017).

As germ cells transition from meiosis I to meiosis II, meiosis I-specific activities must be reversed to allow for sister chromatid segregation during this second equational meiotic division. However, the limited time between the two meiotic stages in many organisms (Kishimoto, 2003) restricts the ability of transcriptional or translational control to broadly rewire the cell division apparatus to distinguish these events. How meiotic cells coordinate this rapid and substantial change to the cell cycle machinery is a critical unanswered question for our understanding of meiosis. Here, we sought to define the mechanisms that enable the switch between meiosis I and II by analyzing the regulatory control of Meikin activity during the different stages of meiosis. Our results demonstrate that Separase cleavage of Meikin acts to rewire cell division activities and create distinct behaviors for each meiotic stage.

## Results

### Meikin is proteolytically cleaved by Separase during anaphase I

To define how germ cells accomplish two distinct cell divisions in rapid succession, it is critical to determine how the activities of meiosis-specific factors, such as Meikin, are precisely controlled. Although Meikin expression is normally restricted to the germline, we found that ectopically-expressed mNeonGreen-tagged Meikin localized to kinetochores during both interphase and mitosis in mitotically-dividing human HeLa cells (Fig. 1A). The ability of Meikin to localize to mitotic kinetochores suggests that its kinetochore binding partners are at least partially retained between meiosis and mitosis, providing an experimentally-tractable system to analyze Meikin behavior. Consistent with the delocalization of Meikin that occurs during anaphase I of meiosis (Kim et al., 2015a), mNeonGreen-Meikin localization was lost from mitotic kinetochores during anaphase (Fig. 1B). The loss of Meikin localization at anaphase onset could reflect changes to its phosphorylation, anaphase-specific degradation, proteolytic-processing, or structural changes to Meikin or its kinetochore binding partners. Interestingly, as cells progressed into anaphase, Western blotting revealed the formation of a faster migrating form of Meikin, suggestive of proteolytic cleavage (Fig. 1C).

**Fig. 1:**
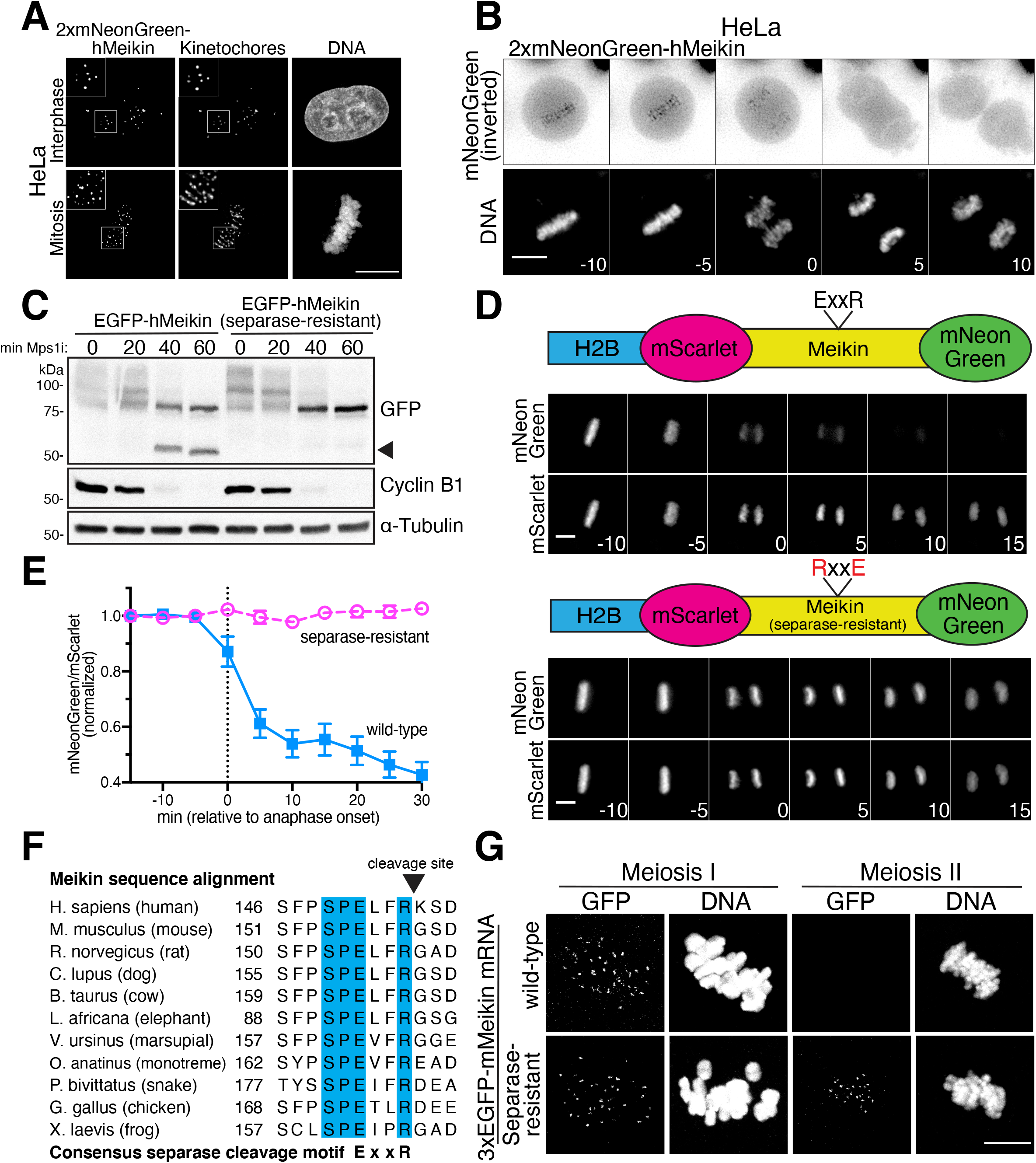
Meikin is cleaved by Separase during anaphase. **A.** Deconvolved immunofluorescence images of HeLa cells stably expressing 2xmNeonGreen-hMeikin. Kinetochores are stained with ACA. Images are not scaled equivalently. Insets, 5 μm. **B.** Montage of deconvolved time-lapse images of HeLa cells expressing 2xmNeonGreen-hMeikin entering anaphase. Numbers indicate minutes relative to anaphase onset. **C.** Western blot showing Meikin mobility during mitotic exit. EGFP-hMeikin expression was induced in HeLa cells by doxycycline treatment. Cells were arrested in mitosis with nocodazole then forced to exit from mitosis by treatment with Mps1i and analyzed at the indicated timepoints. Lower mobility EGFP-hMeikin bands are due to mitosis-specific phosphorylation (see Fig. S3). A high-mobility fragment of EGFP-hMeikin, indicated with the arrow, appears upon mitotic exit. Cyclin B1 degradation indicates mitotic exit. Tubulin was blotted as a loading control. **D.** Schematic of dual-color H2B-cleavage sensors and montage of time-lapse images of HeLa cells expressing the indicated sensor. A Meikin fragment (amino acids 1-332) excluding the kinetochore localization sequence (see Fig. 2) was used. Proteolytic cleavage of Meikin leads to release and diffusion of C-terminal mNeonGreen, but retention of the N-terminal H2B-mScarlet on the DNA. Numbers indicate minutes relative to anaphase onset. **E.** Quantification of Meikin H2B-cleavage sensor. The DNA mass was segmented and the ratio of mNeonGreen:mScarlet within the DNA mass was quantified and normalized to the −15 min timepoint. Error bars indicate 95% confidence interval. 15 cells were analyzed per condition. **F.** Sequence alignment of the Separase cleavage site in Meikin in various vertebrates. Fully conserved amino acids are indicated in blue. Separase is predicted to cleave after the arginine residue in the ExxR motif. **G.** Mouse oocytes injected with the indicated 3xEGFP-mMeikin mRNA were observed in meiosis I and meiosis II for Meikin localization to kinetochores. Images taken at the same timepoint are scaled equivalently. Scale bars, 10 μm. See also Figure S1.

To determine if Meikin is indeed proteolytically cleaved during anaphase, we adapted a previously described protease cleavage sensor (Shindo et al., 2012) in which we targeted Meikin to chromatin through an N-terminal fusion to histone H2B, with fluorescent proteins placed at the Meikin N- and C-termini (Fig. 1D). In this system, Meikin cleavage would result in the delocalization of the mNeonGreen tag, but the chromosomal retention of the H2B-mScarlet tag. Time-lapse imaging revealed that the H2B-mScarlet tagged portion of Meikin remained localized to chromatin as cells exited mitosis. In contrast, the mNeonGreen tagged Meikin fragment progressively delocalized from chromatin during anaphase (Fig. 1D-E). Together, these data indicate that Meikin is proteolytically processed during anaphase when expressed in mitotically-dividing cells.

We next sought to define the protease responsible for Meikin cleavage. Separase is a cysteine protease that cleaves cohesin to initiate sister chromatid separation (Hauf et al., 2001; Uhlmann et al., 1999; Uhlmann et al., 2000). Separase is specifically activated at the metaphase-anaphase transition, and the behavior of the Meikin cleavage sensor mirrors that of a sensor for the established Separase substrate, Rad21/Scc1 (Shindo et al., 2012) (Fig. S1A-B). Indeed, we found that depletion of Separase by RNAi inhibited Meikin proteolysis (Fig. S1C). The consensus cleavage motif for Separase cleavage is ExxR (Hauf et al., 2001; Uhlmann et al., 1999), and charge-swap mutations (<u>R</u>xx<u>E</u>) of this motif in the cohesin subunit Rad21 eliminate its proteolytic processing (Hauf et al., 2001; Shindo et al., 2012) (Fig. S1A-B). Human Meikin contains three <u>E</u>xx<u>R</u> consensus sequences, but only one of these sequences (amino acids 151-154) is conserved across vertebrates (Fig. 1F). We found that charge-swap mutations in this conserved Meikin Separase motif (<u>E</u>LF<u>R</u> to <u>R</u>LF<u>E</u>) eliminated Meikin proteolysis based on analysis of its electrophoretic mobility (Fig. 1C) and our H2B-targeted cleavage sensor (Fig. 1D-E). For some previously established substrates, Separase recognizes an acidic or phosphorylated residue at the P6 position relative to the cleavage site to enhance proteolysis (Alexandru et al., 2001; Hauf et al., 2001). Meikin contains a conserved serine in the P6 position (amino acid 149, Fig. 1F) matching a proline-directed cyclin-dependent kinase (Cdk) phosphorylation motif. Mutation of this residue to alanine reduced Meikin cleavage (Fig. S1D). Together, these data indicate that Meikin is cleaved at anaphase onset by Separase when expressed in mitotic cells.

To determine whether Separase also cleaves Meikin during the meiosis I/II transition in germ cells, we tested the localization of N-terminally tagged murine Meikin in mouse oocytes using mRNA injection. 3xEGFP-Meikin localized to kinetochores in meiosis I, but was lost from kinetochores in meiosis II (Fig. 1G). In contrast, Separase-resistant 3xEGFP-Meikin (E156R, R159E) localized to kinetochores during both meiosis I and meiosis II (Fig. 1G). This suggests that cleavage of Meikin by Separase occurs during anaphase I of meiosis. Together, these data demonstrate that Meikin is processed by the Separase protease at anaphase onset in meiotically dividing cells.

### The Meikin C-terminus is necessary and sufficient for kinetochore localization

Meikin plays a key role in the meiosis I-specific processes of kinetochore co-orientation and sister chromatid cohesion protection (Kim et al., 2015a), critical activities that must be reversed to enable the events associated with meiosis II. Our discovery that Meikin is specifically targeted by Separase during anaphase I provides an attractive model for how Meikin activity is restricted to meiosis I. Thus, we hypothesized that Separase cleavage acts to fully inactivate Meikin, similar to the effect of Separase proteolysis to inactivate cohesin complexes and promote sister chromatid separation (Uhlmann et al., 1999; Uhlmann et al., 2000). To test this hypothesis, we analyzed the consequences of Separase cleavage to Meikin’s known molecular functions. Separase-mediated Meikin cleavage is predicted to produce two protein fragments - N-Meikin (amino acids 1-154) and C-Meikin (amino acids 155-373). Based on the delocalization of the mNeonGreen fluorescence in our cleavage sensor (Fig. 1D), these fragments do not remain associated after proteolysis, but whether these fragments retain any activity is unclear.

To understand how this processing event alters Meikin function at the meiosis I/II transition, we first sought to define the consequences of Meikin cleavage to its kinetochore localization. By analyzing a series of truncation mutants, we identified a Meikin C-terminal domain (amino acids 328-373) that is both necessary and sufficient for kinetochore localization when expressed ectopically in HeLa cells (Fig. 2A). Similarly, we found that the equivalent C-terminal domain of murine Meikin (amino acids 387-434) was sufficient to localize to meiosis I kinetochores in mouse oocytes (Fig. 2B). Previous yeast two-hybrid assays (Kim et al., 2015a) and our affinity purifications of Meikin from mitotic cells (Fig. S2A) both identified interactions with centromere protein C (CENP-C), which is present constitutively at both meiotic and mitotic centromeres (Earnshaw et al., 1989; Kitajima et al., 2011; Tanaka et al., 2009). Meikin also co-localizes with CENP-C to the inner kinetochore (Fig. S2B). Using recombinant proteins, we found that the Meikin C-terminal region was sufficient to bind to CENP-C directly in vitro (Fig. 2C). Mutation of two C-terminal isoleucine-rich motifs (_333_ICCII and _367_DIII) previously implicated in Meikin localization in spermatocytes (Kim et al., 2015a) abolished Meikin’s kinetochore localization in both HeLa cells and mouse oocytes (Fig. 2A-B) and its interaction with CENP-C in vitro (Fig. 2D). In reciprocal experiments, we identified a minimal C-terminal fragment of CENP-C (amino acids 808-943) that is sufficient for Meikin binding (Fig. S2C). As we found that increased salt concentrations promoted Meikin-CENP-C binding (Fig. S2D), we tested whether their interface requires hydrophobic interactions. We identified two hydrophobic patches within CENP-C (_835_IILM or _865_PFF) that are required for Meikin binding (Fig. S2E-F), possibly by partnering with the Meikin isoleucine-rich motifs. This binding site for Meikin on CENP-C does not overlap with established CENP-C-interaction partners (Klare et al., 2015), suggesting that CENP-C interacts with Meikin without disrupting its other kinetochore interfaces.

**Fig. 2:**
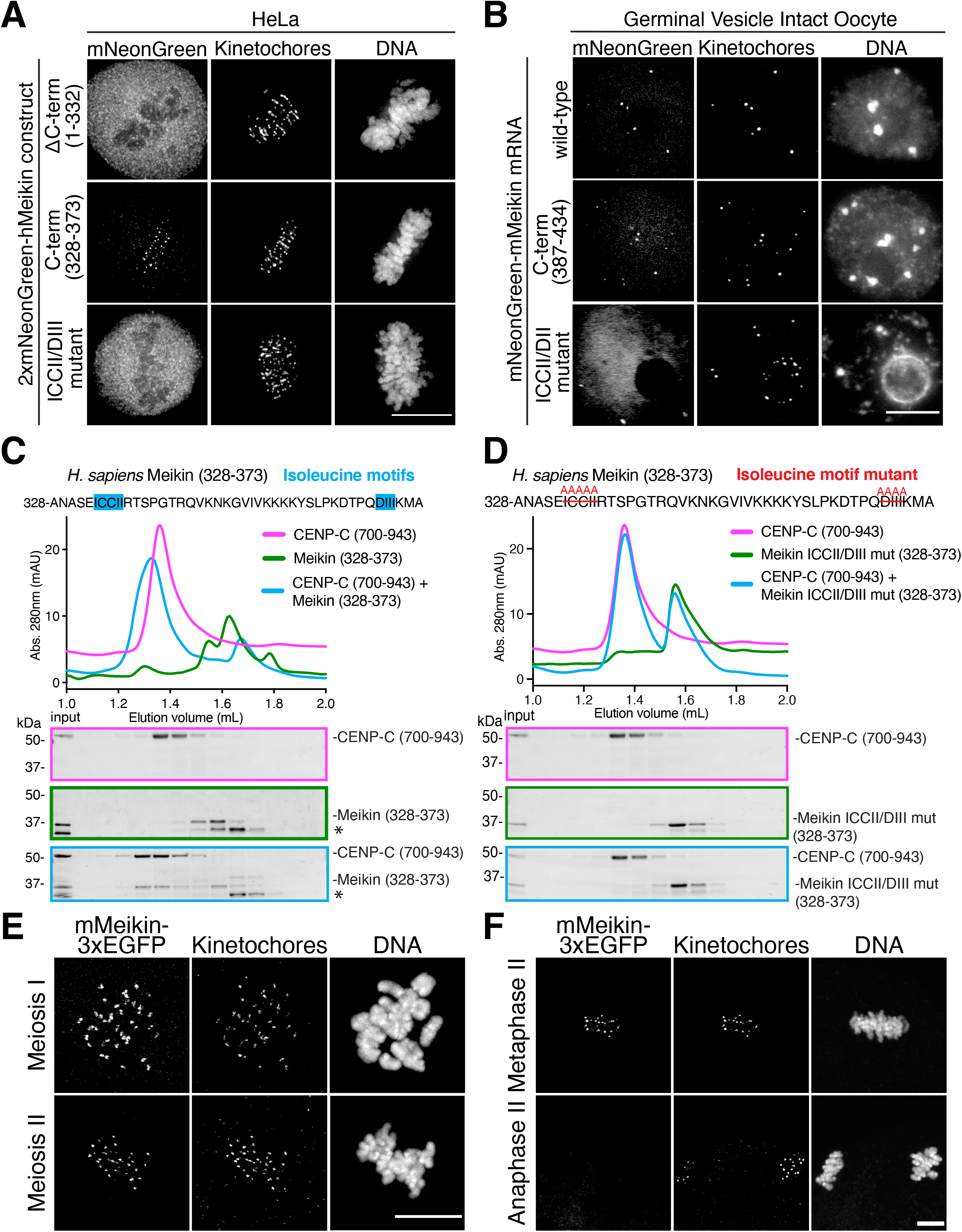
The Meikin C-terminus is necessary and sufficient for kinetochore localization and CENP-C binding. **A.** Deconvolved immunofluorescence images of HeLa cells stably expressing the indicated 2xmNeonGreen-hMeikin constructs. Kinetochores are stained with ACA. Images are not scaled equivalently. **B.** Immunofluorescence images of germinal vesicle intact mouse oocytes injected with the indicated mNeonGreen-mMeikin mRNA. Kinetochores are stained with mouse CENP-A antibody. Images are not scaled equivalently. **C.** Recombinant sfGFP-Meikin and GST-CENP-C protein fragments were bound and complexes analyzed by size exclusion chromatography. Fractions corresponding to elution volumes 1.0 to 2.0 mL were analyzed by SDS-PAGE and Coomassie staining. The Meikin C-terminus containing the isoleucine motifs is sufficient for binding to CENP-C as indicated by co-elution of the two proteins. A contaminating protein in the Meikin prep is indicated with an asterisk. **D.** Recombinant sfGFP-Meikin containing alanine mutations in the isoleucine motifs and GST-CENP-C fragments were analyzed by gel filtration as in Fig. 2C. **E.** Immunofluorescence images of mouse oocytes injected with mMeikin-3xEGFP mRNA and fixed at the indicated stage. Kinetochores are stained with mouse CENP-C antibody. Images are scaled equivalently. **F.** Immunofluorescence images of mouse oocytes injected with mMeikin-3xEGFP mRNA and fixed at the indicated stage. Kinetochores are stained with mouse CENP-C antibody. Images are scaled equivalently. Scale bars, 10 μm. See also Figure S2.

As the Meikin C-terminus is sufficient for kinetochore localization and its interaction partner CENP-C is present throughout meiosis, this suggests that the C-terminal Meikin fragment generated by Separase cleavage would retain the ability to target to kinetochores. Indeed, although N-terminally tagged Meikin in oocytes is lost from kinetochores in meiosis II (Fig. 1G), we found that C-terminally tagged Meikin localizes to kinetochores during both meiosis I and meiosis II (Fig. 2E), and is only removed from kinetochores upon completion of meiosis during anaphase II (Fig. 2F). Thus, while the N-terminal fragment of Meikin is lost from kinetochores following Separase cleavage (Fig. 1G), the Meikin C-terminal cleavage fragment (C-Meikin) remains associated with kinetochores in meiosis II through its interaction with CENP-C, creating the possibility that this fragment could retain some activities at meiosis II kinetochores.

### Plk1 displays phospho-dependent binding to the Meikin C-terminal region

Although Meikin kinetochore localization is retained following Separase cleavage, it is possible that other Meikin interaction partners are affected. Prior work suggested that Meikin promotes meiosis I sister kinetochore co-orientation and the protection of centromeric cohesin through its direct interaction with Polo-like kinase 1 (Plk1) (Kim et al., 2015a), an essential cell cycle kinase that controls multiple aspects of mitosis and meiosis (Petronczki et al., 2008). We found that immunoprecipitation of ectopically-expressed Meikin from HeLa cells arrested in mitosis isolated both CENP-C and Plk1 (Fig. S2A). Plk1-Meikin complexes co-purified from HeLa cells display similar kinase activity on an artificial substrate to Plk1 isolated on its own (Fig. S3A). In addition, ectopic Meikin expression in HeLa cells results in increased Plk1 localization to mitotic kinetochores and increased phosphorylation of Plk1-dependent kinetochore substrates (Fig. S3B-C). The combination of these data suggests that Meikin acts as a targeting subunit for Plk1 to recruit Plk1 to meiotic kinetochores, where it can then phosphorylate downstream targets.

To determine whether Meikin cleavage affects its binding to Plk1, we first sought to define the basis for the Meikin-Plk1 interaction. Plk1 typically binds to its substrates and targeting factors via its phospho-peptide binding domain (Polo-box domain; PBD). We found that Meikin is extensively phosphorylated when expressed in HeLa cells (Fig. 1C; Fig. S3D) on both Cdk ([pT/pS]-P) and Plk1 ([D/E/N]-x-[pT/pS]) consensus phosphorylation sites (Fig. S3E). Chemical inhibition of Plk1 kinase activity eliminates the Meikin-Plk1 interaction (Fig. S4A). Human Meikin contains three S-[pT/pS]-P (STP) motifs (T251, T264, T276), which provide potential docking sites for the Plk1 PBD (Elia et al., 2003), and which have been implicated previously in Plk1 binding (Kim et al., 2015a). Mutation of all three STP motifs (S<u>T</u>P to S<u>A</u>P) reduced, but did not eliminate the Meikin-Plk1 interaction (Fig. 3A). However, a Meikin mutant that eliminates the STP motifs together with multiple consensus sites for Plk1 (S175, T176, T180, S181, S196), which enhance kinase docking interactions in other substrates (Lee et al., 2008), abrogated Plk1 binding (8A mutant; Fig. 3A). Notably, this Meikin-8A mutant is not cleaved efficiently by Separase (Fig. S1D), consistent with other Separase substrates that require phosphorylation for their proteolysis (Alexandru et al., 2001; Hauf et al., 2005; Hornig and Uhlmann, 2004; Kim et al., 2015b; Kudo et al., 2009). We conclude that Meikin binding to Plk1requires its phosphorylation at multiple sites. To determine whether Meikin cleavage affects its binding to Plk1, we next defined the minimal region required for the Meikin-Plk1 interaction. The N-terminal Meikin fragment generated by Separase cleavage (N-Meikin; amino acids 1-154) did not bind to Plk1 (Fig. 3B). In contrast, the C-terminal cleavage fragment (C-Meikin; amino acids 155-373) was still capable of interacting with Plk1 (Fig. 3C). As an alternative strategy to detect Meikin-Plk1 interactions, we targeted Meikin to chromatin using an H2B fusion. Expression of H2B fusions with either full length Meikin or the C-Meikin Separase fragment recruited endogenous Plk1 to chromatin (Fig. S4B). In contrast, an N-Meikin-H2B fusion failed to recruit Plk1 (Fig. S4B). Together, these data suggest that Separase cleavage does not prevent the binding of C-Meikin to Plk1.

**Fig. 3:**
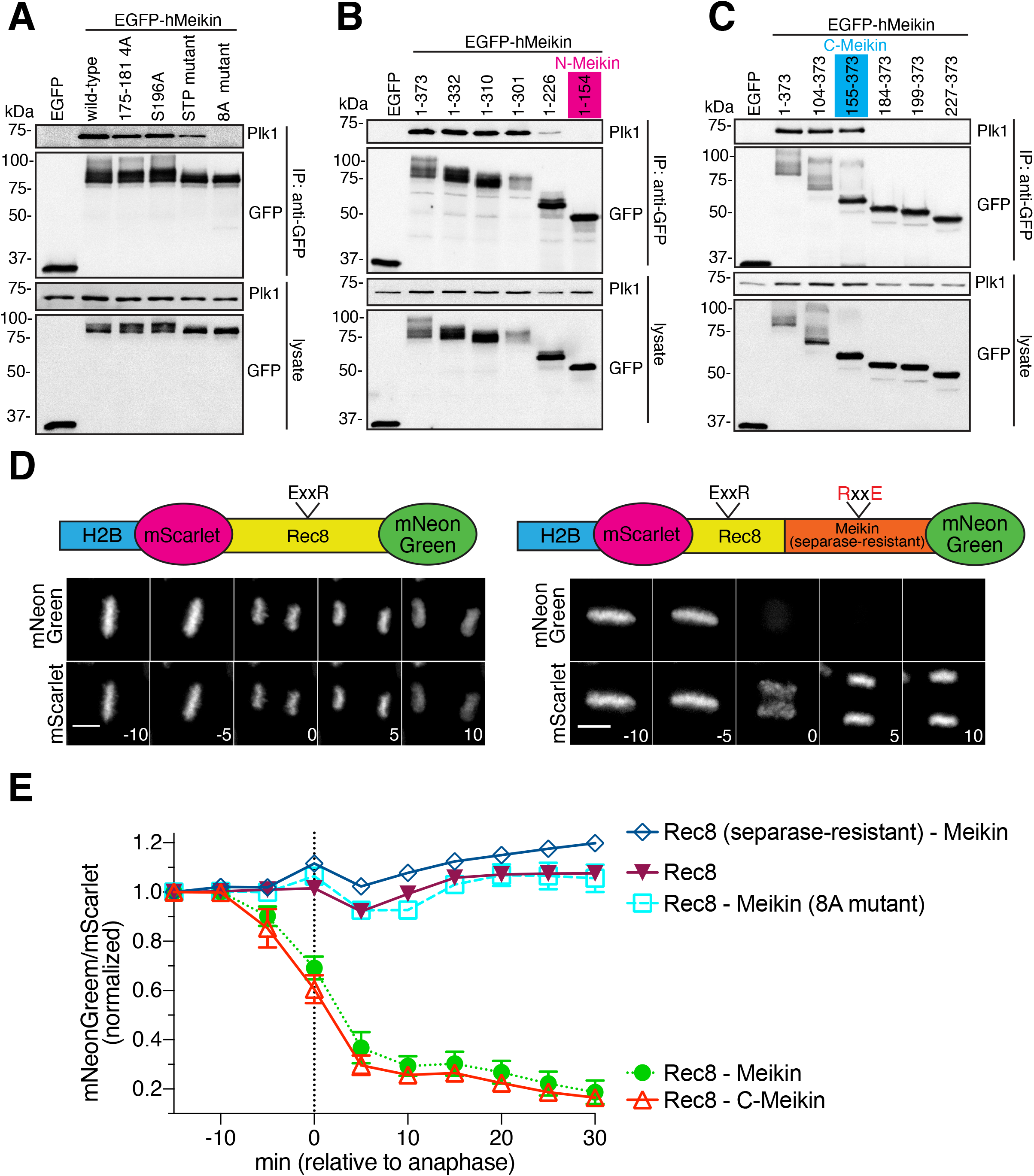
Full lenth Meikin and the C-Meikin cleavage fragment bind Plk1 and promote Rec8 cleavage similarly. **A.** HeLa cells expressing the indicated EGFP-hMeikin C-terminal truncation constructs under the control of a doxycycline inducible promoter were induced and arrested in mitosis by STLC treatment. Cells were lysed and GFP-immunoprecipitation (IP) was performed and analyzed by Western blotting. 175-181 4A mutant includes S175A, T176A, T180A, S181A mutations. STP mutant includes T251A, T264A, T276A mutations. 8A mutant includes S175A, T176A, T180A, S181A, S196A, T251A, T264A, T276A mutations. **B.** Western blot analysis of GFP immunoprecipitates from cells expressing EGFP-hMeikin C-terminal truncation mutants as in Fig. 3A. **C.** Western blot analysis of GFP immunoprecipitates from cells expressing EGFP-hMeikin N-terminal truncation mutants as in Fig. 3A. **D.** Schematic of dual-color Rec8 H2B-cleavage sensors and montage of time-lapse images of HeLa cells expressing the indicated sensor. The sensor contains a fragment of hRec8 (amino acids 297-506) including the predicted Separase cleavage sites and a fragment of hMeikin with Separase-resistant mutations (amino acids 1-332, E151R, R154E). Numbers indicate minutes relative to anaphase onset. Scale bars, 10 μm. **E.** Quantification of Rec8 H2B-cleavage sensor analyzed and represented as in Figure 1. The number of cells analyzed per condition were: Rec8(separase-null)-Meikin (26 cells), Rec8 (30 cells), Rec8-Meikin(8A mutant) (29 cells), Rec8-Meikin (22 cells), Rec8-C-Meikin (22 cells). See also Figure S3 and Figure S4.

### Meikin-Plk1 complexes promote Rec8 cleavage when present in close proximity

We next sought to assess whether Meikin cleavage affects its downstream activities. Current models suggest that Meikin-Plk1 complexes regulate key features of both kinetochore co-orientation and the protection of pericentric cohesin (Kim et al., 2015a). To define the substrates responsible for these activities, we analyzed the meiosis-specific kleisin subunit of cohesin Rec8 (Buonomo et al., 2000; Tachibana-Konwalski et al., 2010). Centromere-localized Rec8 has been proposed to play a critical role in promoting kinetochore co-orientation in many organisms (Chelysheva et al., 2005; Sakuno et al., 2009; Severson et al., 2009; Watanabe and Nurse, 1999). In this model, the centromeric Rec8 population would need to be eliminated at the meiosis I/II transition to allow for proper chromosome alignment at metaphase II. In contrast, the pericentric population of Rec8-containing cohesin must be protected from Separase cleavage to ensure that sister chromatids remain associated until anaphase II. Indeed, recent work suggests that the centromeric, pericentric, and chromosome arm populations of Rec8 are differentially regulated at the meiosis I/II transition in mammalian oocytes (Ogushi et al., 2020).

To monitor Rec8 cleavage, we adapted the H2B-based cleavage sensor in HeLa cells. In contrast to the Rad21 cleavage sensor (Fig. S1B), a Rec8 fragment containing its established Separase cleavage sites did not show any proteolysis upon anaphase onset in mitotic cells (Fig. 3D-E). To test whether Meikin-Plk1 complexes could promote Rec8 cleavage when present in close proximity, we created an in-frame Rec8-Meikin fusion. Strikingly, fusion of Meikin to the Rec8 fragment resulted in efficient Rec8 cleavage (Fig. 3D-E). Charge-swap mutation of the conserved Separase cleavage site in Rec8 (E401R, R404E) (Kudo et al., 2009) eliminated this proteolysis (Fig. 3E). Rec8 cleavage was also dependent on the interaction between Meikin and Plk1, as a Meikin mutant that does not bind Plk1 (8A mutant; Fig. 3A) does not potentiate Rec8 cleavage (Fig. 3E). This suggests that kinetochore-targeted Meikin-Plk1 complexes could phosphorylate Rec8 to promote the cleavage of adjacent centromere-proximal Rec8-cohesin at anaphase I. Consistent with this, prior work found that Plk1 phosphorylation of mammalian Rec8 enhances Separase-mediated cleavage in vitro (Kudo et al., 2009). These data suggest that Meikin-Plk1 complexes activate proximal Rec8 for cleavage by Separase, providing a mechanism to reverse kinetochore co-orientation following anaphase I.

To determine whether Separase-cleavage inhibits Meikin’s ability to promote Rec8 cleavage, we next created an in-frame fusion between the C-Meikin cleavage product and Rec8 in our H2B sensor. Similarly to full length Meikin, C-Meikin induced efficient cleavage of the Rec8 sensor (Fig. 3E). This suggests that retention of C-Meikin at kinetochores would further promote the elimination of centromeric Rec8 during late anaphase, ensuring complete reversal of kinetochore co-orientation before meiosis II chromosome alignment. Importantly, these data indicate that full length Meikin and the C-Meikin Separase cleavage product display similar activities with respect to their kinetochore localization, Plk1 binding, and ability to promote Rec8 cleavage. Thus, in contrast to other substrates, Separase cleavage does not fully inactivate Meikin function.

### A sensitized assay for Plk1 interactions reveals distinct behaviors for full-length and C-Meikin

Given the similar activities observed for full-length Meikin and C-Meikin in the assays described above, we sought to generate a sensitized assay for Meikin activity. Stable expression of Meikin at low levels in mitotically dividing cells results in a modest increase in misaligned chromosomes (Fig. S5A). In contrast, increased Meikin expression (under the control of a doxycycline-inducible promoter), caused a potent mitotic arrest with misaligned chromosomes and monopolar spindle structures (Fig. S5A-B), phenotypes consistent with an effect on Plk1 activity. The ability of Meikin overexpression to disrupt cell division provides a sensitized assay to monitor the behaviors of Meikin mutants. For example, although individual Meikin phosphorylation site mutants were not sufficient to disrupt Meikin-Plk1 binding (Fig. 3A), these mutants did not promote a mitotic arrest (Fig. S5C). This suggests that a direct interaction between Meikin and Plk1 is required, but not sufficient for the mitotic arrest phenotype. Utilizing this assay, we defined a minimal domain (amino acids 124-332) spanning the N- and C-Meikin Separase cleavage fragments that is sufficient to induce a mitotic arrest (Fig. S5C). Importantly, neither the N-nor C-terminal Separase cleavage fragment of Meikin was sufficient to generate a mitotic arrest (Fig. S5C), despite an interaction between C-Meikin and Plk1 (Fig. 3C). Based on these data, we conclude that full-length Meikin and C-Meikin both localize to kinetochores and bind to Plk1, but display different efficiency or properties for their Plk1 interactions. Thus, Separase cleavage has the potential to differentially control Meikin-Plk1 interactions at the meiosis I/II transition.

### Separase cleavage of Meikin is required for chromosome alignment during meiosis II

Based on the results described above, we sought to directly test whether Separase-mediated Meikin proteolysis is required for the proper execution of meiosis. To determine the functional consequences of Meikin cleavage, we ectopically expressed the Separase-resistant Meikin mutant in mouse oocytes, causing full length Meikin to persist at kinetochores into meiosis II (Fig. 1G; Fig. 4A). Despite the presence of endogenous Meikin, the induced retention of Meikin at kinetochores during meiosis II in this Separase cleavage mutant caused severe chromosome alignment defects during the second meiotic division (Fig. 4B-C). Separase-resistant Meikin expression also resulted in increased kinetochore levels of Plk1 and Bub1 at meiosis II kinetochores (Fig. S5D-E). As Plk1 and Bub1 regulate kinetochore-microtubule interactions (Petronczki et al., 2008; Watanabe, 2012), improper kinetochore localization of Plk1 and Bub1 in meiosis II oocytes expressing Separase-resistant Meikin likely contributes to the observed chromosome alignment defects (Fig. 4B-C). Thus, Meikin cleavage during anaphase I is critical to enable a proper meiosis II division.

**Fig. 4:**
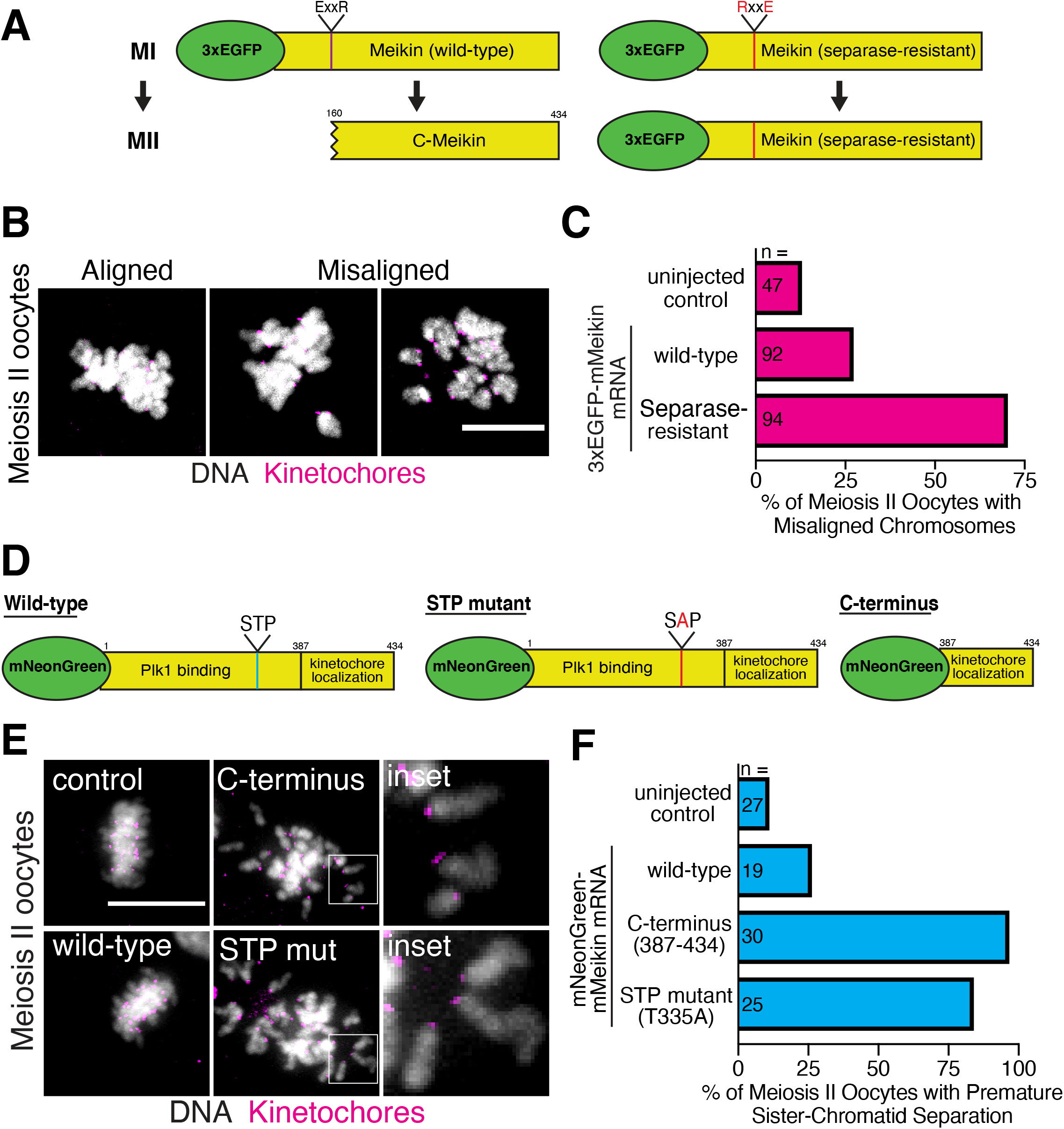
Separase cleavage of Meikin is required for proper meiosis II chromosome alignment. **A.** Schematic of murine Meikin constructs injected and their expected behaviors at meiosis II. Separase-resistant Meikin contains a charge swap mutation in the Separase cleavage site (E156R, R159E). **B.** Representative immunofluorescence images of chromosome misalignment defects observed in meiosis II oocytes. **C.** Quantification of meiosis II chromosome misalignment defects observed in oocytes injected with the indicated 3xEGFP-mMeikin construct. **D.** Schematic of murine Meikin constructs injected. **E.** Representative immunofluorescence images of mouse oocytes injected with the indicated mNeonGreen-mMeikin mRNA and matured to meiosis II. Insets show separated chromatids. **F.** Quantification of premature sister chromatid separation defects observed in meiosis II oocytes expressing the indicated Meikin mRNA. n represents the number of meiosis II oocytes analyzed. Kinetochores are stained with Hec1 antibody. Scale bars, 10 μm. Insets, 5 μm. See also Figure S5.

### C-Meikin is required during meiosis II for proper chromosome alignment

Our results indicate that the failure to cleave Meikin results in a defective meiosis II, but that cleaved C-Meikin retains at least partial activity based on its kinetochore localization, Plk1 binding, and ability to promote Rec8 cleavage. Thus, instead of abolishing Meikin function, our work suggests that Separase processing modulates Meikin activity at anaphase I to promote additional functions during meiosis II. To test this, we sought to disrupt Meikin function by creating dominant-negative Meikin alleles. For these experiments, we first expressed Meikin mutants that retain its CENP-C interaction (allowing them to displace endogenous Meikin at kinetochores), but are defective for Plk1 binding (Fig. 4D). Oocytes ectopically expressing wild-type Meikin using mRNA microinjection progressed to metaphase II similarly to mock-injected oocytes. In contrast, expression of the minimal C-terminal murine Meikin kinetochore targeting domain (amino acids 387-434; corresponding to amino acids 328-373 in human Meikin), which lacks Plk1 binding (Fig. 3C), caused premature-sister chromatid separation during meiosis II (Fig. 4E-F). Similarly, expression of full length Meikin with a mutation in the STP motif also caused premature-sister chromatid separation during meiosis II (Fig. 4E-F). These phenotypes suggest that the Meikin-Plk1 interaction acts during meiosis I to protect the pericentric cohesion that holds sister chromatids together until anaphase II.

To test Meikin function during meiosis II, we next developed a conditional strategy to fully eliminate Meikin activity at anaphase I onset, allowing us to circumvent the consequences of the dominant negative Meikin mutants that disrupt meiotic behaviors in meiosis I (Fig. 4D-F). To do this, we altered the position of Separase proteolysis by inserting the Rad21 Separase cleavage site between the Meikin Plk1 binding and kinetochore targeting domains defined by our analysis (Fig. 2A-B, Fig. 3B-C), forming a Meikin allele with two potential cleavage sites (Fig. 5A). Introducing inactivating mutations at either the endogenous Meikin cleavage site or the inserted cleavage site enabled us to control the position of Separase proteolysis to generate either a product equivalent to C-Meikin or a product completely lacking Plk1 binding activity. Importantly, both alleles retain full Meikin activity during meiosis I, allowing us to determine the functional consequences of specifically inactivating Meikin during meiosis II. Expression of the control Meikin construct which is cleaved at the endogenous Meikin proteolysis site (amino acid 160) resulted in normal meiosis II oocytes (Fig 5B-C). In contrast, targeting Separase to the introduced downstream site (amino acid 387) to eliminate Meikin-Plk1 interactions during meiosis II resulted in chromosome alignment defects in meiosis II oocytes (Fig 5B-C). These data indicate that C-Meikin is not simply an inactive form of Meikin, but is required for proper chromosome alignment after anaphase I. Thus, instead of Meikin acting as a meiosis I-specific factor as proposed by prior work (Kim et al., 2015a), Meikin remains active throughout meiosis with Separase cleavage at anaphase I generating a distinct form of Meikin with different activity. Full length Meikin acts to enable meiosis I events, such as kinetochore co-orientation and the protection of pericentric cohesin (Kim et al., 2015a), with the C-Meikin cleavage product retaining modified activities that are critical to enable meiosis II events (Fig. 5D).

**Fig 5:**
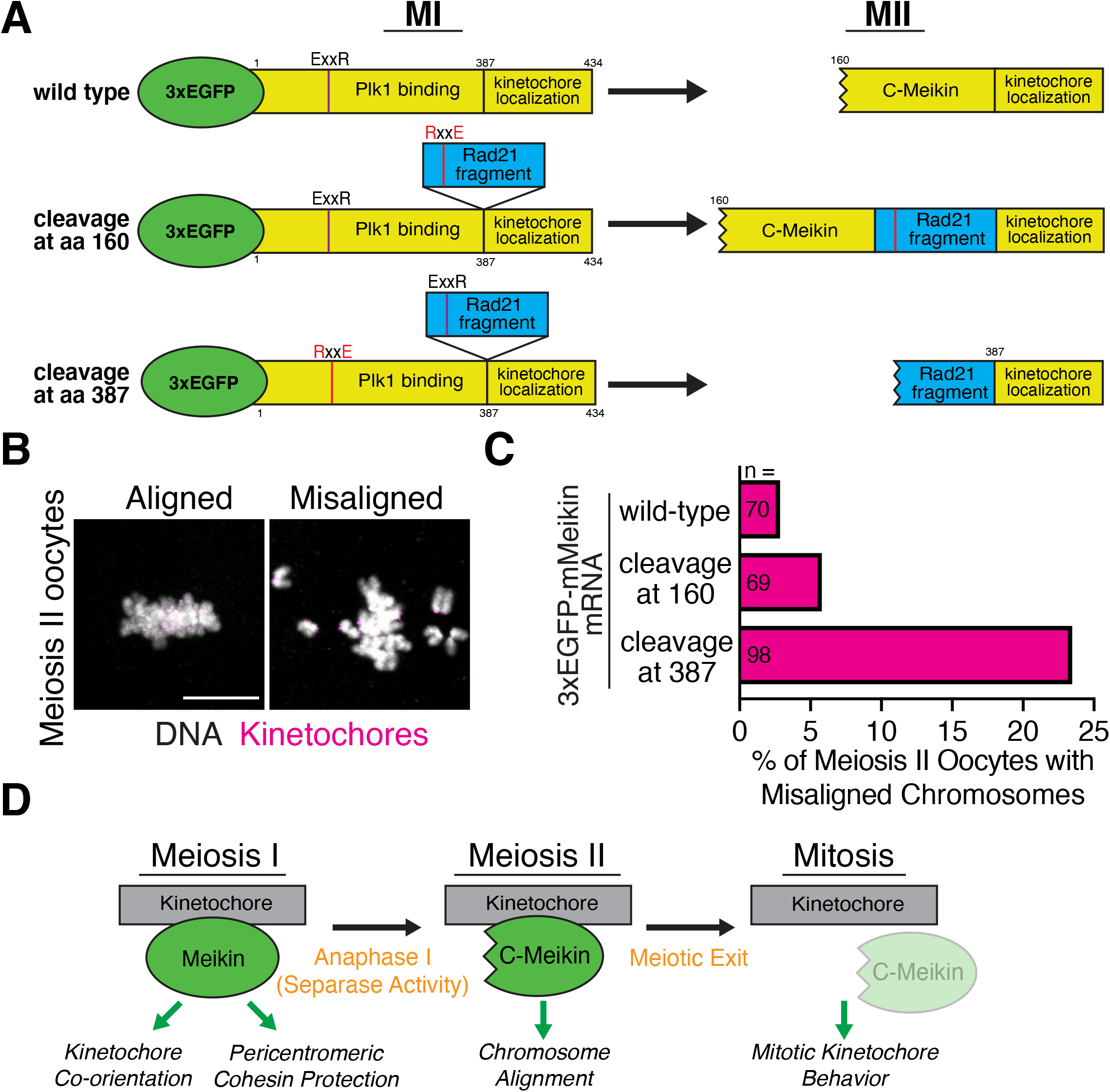
The C-Meikin cleavage fragment is required for meiosis II chromosome alignment. **A.** Schematic of murine Meikin constructs injected and their expected behaviors at meiosis II. A fragment of hRad21 (amino acids 142-275) was inserted between the Plk1 binding and kinetochore localization domains of Meikin. Charge-swap mutations in the Separase-cleavage sites of Meikin (E156R, R159E) or Rad21 (E169R, R172E) were used to direct the location of Separase cleavage during anaphase I. **B.** Representative immunofluorescence images of chromosome misalignment defects observed in meiosis II oocytes. Kinetochores are stained with mouse CENP-C antibody. Scale bars, 10 μm. **C.** Quantification of meiosis II chromosome misalignment defects observed in oocytes injected with the indicated 3xEGFP-mMeikin construct. n represents the number of meiosis II oocytes analyzed. **D.** Model for differential Meikin activity at distinct stages of meiosis.

## DISCUSSION

Our data reveal the intricate regulatory control by which the two meiotic divisions are coordinated, providing a mechanism to rewire the cell division machinery between meiosis I and II. We find that the protease Separase acts as a molecular “scalpel” to precisely and irreversibly modulate Meikin activity between the two meiotic divisions. Instead of proteolytic cleavage completely inactivating the protein, full-length Meikin and the C-Meikin Separase cleavage product both localize to kinetochores and bind Plk1, but differ in their functional activities. Importantly, the activity of C-Meikin during meiosis II is required for proper chromosome alignment and cannot be replaced with full-length Meikin. Both the failure to cleave Meikin (Fig. 4A-C) or the complete inactivation of Meikin at anaphase I (Fig. 5A-C) result in defective chromosome alignment during meiosis II. Thus, Separase cleavage of Meikin is critical for mammalian meiosis. Notably, Meikin counterparts in other organisms have distinct mechanisms of regulation that restrict their activities to meiosis I. For example, the budding yeast meiosis I factor Spo13 is degraded at anaphase I by the APC/C (Sullivan and Morgan, 2007), which results in its elimination.

Our data support a model in which kinetochore-localized Meikin establishes a gradient of Plk1 activity during meiosis I centered on the kinetochore, which leads to different fates for cohesin complexes based on their location within the gradient. Meikin is targeted to kinetochores through its binding to CENP-C (Fig. 2A-D), allowing Meikin to recruit Plk1 (Fig. S3C-D). Centromere-localized Meikin-Plk1 complexes differentially regulate the centromeric and pericentromeric populations of Rec8 cohesin during meiosis, with opposing effects on these two pools. By increasing the concentration of centromere-localized Plk1, Meikin-Plk1 complexes direct phosphorylation of centromere-bound Rec8 to promote its efficient cleavage during anaphase I (Fig. 3D-E), thereby reversing meiosis I kinetochore co-orientation. In parallel, Meikin-Plk1 activity recruits Bub1 (Fig. S5D-E), which in turn recruits Sgo2-PP2A complexes to the pericentromere to reverse Rec8 phosphorylation and protect pericentromeric Rec8 from Separase cleavage (Fig. 4E-F) (Galander et al., 2019a; Galander et al., 2019b; Marston, 2015; Miyazaki et al., 2017). Thus, during anaphase I, Separase cleaves centromere-proximal and phosphorylation-primed Rec8, but not pericentromeric Rec8, eliminating kinetochore co-orientation while maintaining sister chromatid cohesion. At anaphase I, Separase also cleaves Meikin (Fig. 1G), resulting in a C-terminal Meikin fragment that remains associated with kinetochores (Fig. 2E). However, while retaining key activities, this cleaved version is less potent in its ability to bind to or regulate Plk1 (Fig. S5B-C), leading to reduced Plk1 at kinetochores in meiosis II compared to meiosis I (Fig. S5D-E) (Kim et al., 2015a). This reduced Plk1 activity leads to a corresponding reduction in pericentromeric Bub1 and Sgo2, allowing for Rec8 phosphorylation and cleavage at anaphase II. This model is consistent with recent findings that Separase activity at meiosis I is required for deprotection of peri-centromeric cohesin (Gryaznova et al., 2020; Ogushi et al., 2020). Finally, as germ cells exit meiosis, Meikin localization is completely eliminated (Fig. 2F), returning kinetochore regulation to a mitotic state.

In conclusion, Meikin is a novel Separase substrate that is cleaved specifically during anaphase of meiosis I. Despite the identification of Separase more than two decades ago and extensive efforts to identify additional targets, the number of confirmed Separase substrates remains low. Our data show that Meikin cleavage by Separase in anaphase I does not inactivate the protein, but rather modulates its activity to allow Meikin to transition from meiosis I to meiosis II specific functions. This provides an elegant mechanism to rapidly reverse the meiosis I-specific modifications to the cell cycle machinery and coordinate the sequential meiotic divisions.

## Acknowledgements

We thank the members of the Cheeseman and Lampson labs for their support and input. This work was supported by grants from The Harold G & Leila Y. Mathers Charitable Foundation to IMC, the NIH/National Institute of General Medical Sciences (R35GM126930 to IMC and GM122475 to MAL), and the Henry and Frances Keany Rickard Fund Fellowship from the Massachusetts Institute of Technology Office of Graduate Education to NKM.

## Author Contributions

Conceptualization – NKM, IMC, MAL; Methodology – NKM, JM; Validation – NKM, JM; Investigation – NKM, JM; Writing - Original Draft Preparation – NKM, IMC; Writing – Review & Editing – NKM, IMC, MAL; Visualization – NKM, JM; Supervision: IMC, MAL; Funding Acquisition: IMC, NKM, MAL

## Methods

### Resource Availability

#### Lead contact

All requests for data and materials should be addressed to Iain Cheeseman.

#### Materials Availability

All plasmids and cell lines generated in this study are available upon request from the lead contact.

#### Data and Code Availability

The datasets and code used in this study and not included in the text are available upon request from the lead contact.

### Experimental Model and Subject Details

#### Cell Culture

HeLa cells (transformed human female cervical epithelium) were cultured in Dulbecco’s modified Eagle medium supplemented with 10% fetal bovine serum, 100 U/mL penicillin and streptomycin, and 2 mM L-glutamine at 37°C with 5% CO_2_. Doxycycline inducible cell lines were cultured in medium containing FBS certified as tetracycline free and were induced by addition of doxycycline to 1 μg/mL for 16 hr. Other drugs used on human cells were kinesin Eg5 inhibitor (S-trityl-L-cysteine, STLC, 10 μM), nocodazole (0.3 μM), AZ-3146 (MPS1i, 3 μM), BI-2536 (Plk1i, 10 μM), RO-3306 (Cdk1i, 10 μM). Hela cells were regularly monitored for mycoplasma contamination using commercial detection kits.

#### Mouse oocyte collection and culture

8-14 week-old female mice used in this study were purchased from Envigo (Strain: NSA(CF-1)). All animal experiments were approved by the University of Pennsylvania Institutional Animal Care and Use Committee and were consistent with the National Institutes of Health guidelines. Female mice were hormonally primed with 5U of Pregnant Mare Serum Gonadotropin (PMSG, Calbiochem, cat# 367222) 44-48 hr prior to oocyte collection. Germinal vesicle intact oocytes were collected in M2 medium (Sigma, M7167), denuded from cumulus cells, and cultured in CZB medium (Sigma, MR-019-D) covered with mineral oil (Sigma, M5310) in a humidified atmosphere of 5% CO_2_ in air at 37°C. During collection, meiotic resumption was inhibited by addition of 2.5 μM milrinone (Sigma, M4659).

### Method Details

#### Cell line generation

The cell lines used in this study are described in Table S2. pBABE derivatives were transfected with Effectene (Qiagen) according to the manufacturer’s protocol along with VSVG packaging plasmid into 293-GP cells for generation of retrovirus as described(Morgenstern and Land, 1990). Supernatant-containing retrovirus was sterile filtered, supplemented with 20 μg/mL polybrene (Millipore) and used to transduce HeLa cells. Doxycycline-inducible cell lines were generated by homology-directed insertion into the AAVS1 “safe-harbor” locus. Donor plasmid containing selection marker, the tetracycline-responsive promoter, the transgene, and reverse tetracycline-controlled transactivator flanked by AAVS1 homology arms(Qian et al., 2014) was transfected using Effectene with a pX330-based plasmid(Cong et al., 2013) expressing both spCas9 and a guide RNA specific for the AAVS1 locus (pNM220, gRNA sequence – 5’-GGGGCCACTAGGGACAGGAT). Two days post-transfection or transduction, cells were selected with the appropriate antibiotic (puromycin at 0.5 μg/mL or blasticidin at 2 μg/mL, Life Technologies). Where indicated, clonal lines were obtained by fluorescence activated cell-sorting single cells into 96 well plates. Fluorescence enriched lines were generated by bulk sorting a polyclonal population for fluorescence positive cells.

#### Immunofluorescence and microscopy of mitotic cells

Cells for immunofluorescence were seeded on poly-L-lysine (Sigma-Aldrich) coated coverslips and fixed in PBS plus 4% formaldehyde for 10 min unless otherwise noted in figure legends. Coverslips were washed with PBS plus 0.1% Triton X-100 and blocked in Abdil (20 mM Tris-HCl, 150 mM NaCl, 0.1% Triton X-100, 3% bovine serum albumin, 0.1% NaN_3_, pH 7.5). Primary antibodies used in this study are described in Table S1 and were diluted in Abdil. Cy3- and Cy5-conjugated secondary antibodies (Jackson ImmunoResearch Laboratories) were diluted 1:300 in PBS plus 0.1% Triton X-100. DNA was stained with 1 μg/mL Hoechst-33342 (Sigma-Aldrich) in PBS plus 0.1% Triton X-100 for 10 min. Coverslips were mounted using PPDM (0.5% *p*-phenylenediamine, 20 mM Tris-HCl, pH 8.8, 90% glycerol). Images were acquired on a DeltaVision Core deconvolution microscope (Applied Precision) equipped with a CoolSnap HQ2 charge-coupled device camera and deconvolved where appropriate. All images are maximal projections in z unless otherwise indicated. Image analysis was performed in Fiji (ImageJ, NIH) (Schindelin et al., 2012).

Integrated fluorescence intensity of mitotic kinetochores was measured with a custom CellProfiler pipeline (McQuin et al., 2018). The median intensity of a 5-pixel wide region surrounding each kinetochore was used to background subtract each measurement.

#### Live-cell imaging of mitotic cells

For live-cell imaging, cells were seeded into 8-well glass-bottomed chambers (Ibidi) and moved into CO_2_-independent media (Life Technologies) before imaging at 37°C. For certain movies, DNA was stained with SiR-DNA (Cytoskeleton Inc) at 0.2 μM. Images were acquired on a DeltaVision Core deconvolution microscope (Applied Precision) equipped with a CoolSnap HQ2 charge-coupled device camera and deconvolved where appropriate. Image analysis was performed in Fiji (ImageJ, NIH). For quantification of H2B cleavage sensors, cells undergoing anaphase were selected in Fiji then analyzed using a custom CellProfiler pipeline (McQuin et al., 2018). The pipeline segmented cells based on SiR-DNA signal, performed background subtraction, and measured mNeonGreen and mScarlet signal in each cell. The mNeonGreen/mScarlet ratio was normalized to the first timepoint (t = −15 min).

#### RNAi treatment

siRNAs against ESPL1 (5’-GCUUGUGAUGCCAUCCUGAUU) (Waizenegger et al., 2002) and non-targeting control pool (D-001810-10) were obtained from Dharmacon. 5 μL of 20 μM stock siRNA was mixed with 5 μL of Lipofectamine RNAiMax (Life Technologies) and diluted in 500 μL of OptiMem (Life Technologies). The reaction was applied to cells in a 6-well dish. Transfection media was changed after 24 hr.

#### Mitotic index determination

Meikin expressing cells were induced with 1 μg/mL doxycycline for 24 hr. Cells were collected by incubation for 10 min in PBS + 5 mM EDTA, washed once in PBS, then fixed in PBS + 2% formaldehyde for 10 min at room temperature. Cells were blocked in Abdil for 30 min followed by immunostaining for phosphorylated S10 on histone 3 followed by Cy-5 conjugated secondary antibody. The proportion of GFP-positive single cells also staining positive for H3pS10 was determined on an LSRFortessa (BD Biosciences) flow cytometer and analyzed with FACSDiva software (BD Biosciences). Over 5,000 GFP-positive cells were analyzed per condition.

#### GFP immunoprecipitation and Mass-spectrometry

IP-MS experiments were performed as described previously (Cheeseman and Desai, 2005). Harvested cells were washed in PBS and resuspended 1:1 in 1X Lysis Buffer (50 mM HEPES, 1 mM EGTA, 1 mM MgCl_2_, 100 mM KCl, 10% glycerol, pH 7.4) then drop frozen in liquid nitrogen. Cells were thawed after addition of an equal volume of 1.5X lysis buffer supplemented with 0.075% Nonidet P-40, 1X Complete EDTA-free protease inhibitor cocktail (Roche), 1 mM phenylmethylsulfonyl fluoride, 20 mM beta-glycerophosphate, 1 mM sodium fluoride, and 0.4 mM sodium orthovanadate. Cells were lysed by sonication and cleared by centrifugation. The supernatant was mixed with Protein A beads coupled to rabbit anti-GFP antibodies (Cheeseman lab) and rotated at 4°C for 1 hr. Beads were washed five times in Wash Buffer (50 mM HEPES, 1 mM EGTA, 1 mM MgCl_2_, 300 mM KCl, 10% glycerol, 0.05% NP-40, 1 mM dithiothreitol, 10 μg/mL leupeptin/pepstatin/chymostatin, pH 7.4). After a final wash in Wash Buffer without detergent, bound protein was eluted with 100 mM glycine pH 2.6. Eluted proteins were precipitated by addition of 1/5^th^ volume trichloroacetic acid at 4°C overnight. Precipitated proteins were reduced with TCEP, alkylated with iodoacetamide, and digested with mass-spectrometry grade Lys-C and trypsin (Promega). Digested peptides were cleaned up using C18 spin columns (Pierce) according to the manufacturer’s instructions. Samples were analyzed on an LTQ XL Ion Trap mass spectrometer (Thermo Fisher) coupled with a reverse phase gradient over C18 resin. Data were analyzed using SEQUEST.

#### GFP immunoprecipitation and Western blot

For IP-Western experiments, cells were harvested, washed once in PBS, then lysed on ice for 15 min in Lysis Buffer (50 mM HEPES, 1 mM EGTA, 1 mM MgCl_2_, 100 mM KCl, 10% glycerol, 1% Triton X-100, 0.05% NP-40, 1 mM dithiothreitol, 1X Complete EDTA-free protease inhibitor cocktail (Roche), 1 mM phenylmethylsulfonyl fluoride, 20 mM β-glycerophosphate, 1 mM sodium fluoride, and 0.4 mM sodium orthovanadate, pH 7.4). Cellular debris was removed by centrifugation. Protein concentrations in each sample were measured using Bradford reagent (Bio-Rad), and sample concentrations were normalized before addition of Protein A beads (Bio-Rad) coupled to affinity-purified rabbit anti-GFP polyclonal antibodies (Cheeseman lab). After 1 hr incubation at 4°C, beads were washed 3X with Wash Buffer (50 mM HEPES, 1 mM EGTA, 1 mM MgCl_2_, 100 mM KCl, 10% glycerol, 0.05% NP-40, 1 mM dithiothreitol, 10 μg/mL leupeptin/pepstatin/chymostatin, pH 7.4). Beads were then incubated in an equal volume of Laemmli buffer for 5 min at 95°C to remove bound protein. Samples were analyzed by SDS-PAGE and Western blotting. Samples were separated by SDS-PAGE and semidry transferred to nitrocellulose. Membranes were blocked for 1h in Blocking Buffer (5% milk in TBS + 0.1% Tween-20). Primary antibodies were diluted in Blocking Buffer + 0.2% NaN_3_ and applied to the membrane for 1 hr. HRP-conjugated secondary antibodies (GE Healthcare) were diluted 1:10,000 in TBS + 0.1% Tween-20 and applied to the membrane for 1 hr. After washing in TBS + 0.1% Tween-20, Clarity enhanced chemiluminescence substrate (Bio-Rad) was added to the membrane according to the manufacturer’s instructions. Membranes were imaged with a KwikQuant Imager (Kindle Biosciences). Membranes were stripped (55°C, 1 hr) in stripping buffer (60 mM Tris-HCl, pH 6.8, 2% SDS, 100 mM β-mercaptoethanol) and re-blocked before re-probing.

#### Anaphase synchronization and cell lysis

For synchronization in anaphase, cells were arrested by treatment with STLC for 16 hr followed by addition of Mps1i for the times indicated in the figure legends. Cells were harvested and washed once in PBS. Cell pellets were resuspended in radioimmunoprecipitation buffer (RIPA, ThermoFisher) supplemented with 1X Complete EDTA-free protease inhibitor cocktail (Roche), 1 mM phenylmethylsulfonyl fluoride, 20 mM β-glycerophosphate, 1 mM sodium fluoride, and 0.4 mM sodium orthovanadate. Cells were lysed on ice for 15 min, and cellular debris was removed by centrifugation at >10,000 g for 10 min at 4°C. Protein concentrations were measured and normalized using a bicinchoninic protein assay (Pierce). Samples were analyzed by SDS-PAGE and Western blot.

#### Phosphatase treatment of lysates

Cells treated with the drugs indicated in the Fig. legends were collected, washed once with PBS then lysed in Lysis Buffer without phosphatase inhibitors (50 mM HEPES, 1 mM EGTA, 1 mM MgCl_2_, 100 mM KCl, 10% glycerol, 0.05% NP-40, 1 mM dithiothreitol, 1X Complete EDTA-free protease inhibitor cocktail (Roche), 1 mM phenylmethylsulfonyl fluoride, pH 7.4) for 10 min on ice. Cellular debris was removed by centrifugation and protein concentrations were measured by Bradford assay (Bio-Rad) and normalized. Cell lysates were supplemented with 1X Protein MetalloPhosphatase buffer (New England Biolabs) and 1 mM MnCl_2_. 1 μL of Lambda Protein Phosphatase (New England Biolabs) or 1 μL of phosphatase inhibitor mix (20 mM β-glycerophosphate, 1 mM sodium fluoride, and 0.4 mM sodium orthovanadate) was added to each 50 μL reaction. After incubation at 30°C for 30 min, reactions were stopped by addition of 2X Laemmli buffer. Samples were analyzed by SDS-PAGE and Western blot.

#### In vitro Plk1 activity assays

Plk1 and Plk1/Meikin complexes were immunoprecipitated from HeLa cells. Cells were induced to express EGFP-Plk1 or EGFP-Meikin by treatment with doxycycline and arrested in mitosis by STLC for 16 hr. Cells were harvested, lysed and GFP-immunoprecipitated as above for mass-spectrometry. Bound proteins were eluted by addition of recombinant Tobacco Etch Virus (TEV) protease (Cheeseman lab) and incubation at 4°C for 16 hr which leads to cleavage between EGFP and the tagged-protein. The amount of Plk1 in each elution was normalized by Western blot.

Enzyme kinetic parameters were measured in a continuous read fluorescence assay (PhosphoSens CSKS-AQT0691K, Assay Quant) according to manufacturer directions in a SpectraMax iD3 plate reader (Molecular Devices) in 96-well format at 30°C with reads every 60 sec. Substrate concentrations were varied from 1-35 μM. Initial reaction rates were fit to the Michaelis-Menton equation in GraphPad Prism.

#### Recombinant protein expression and purification

Plasmids used for recombinant protein expression were based on the pGEX-6P1 backbone and are described in Table S3. BL21(DE3) LOBSTR cells (Andersen et al., 2013) carrying the pRARE tRNA plasmid were transformed with the appropriate plasmid and plated on Luria-Bertani (LB)-agar plates containing the appropriate antibiotic. Overnight liquid cultures of LB supplemented with antibiotics and 4% glucose were grown overnight at 37°C from single colonies. The saturated overnight culture was diluted 1:100 and grown to an OD600nm of 0.6-0.7 at 30°C. Cells were shifted to 16°C and induced with 0.3 mM isopropyl β-D-1-thiogalactopyranoside and incubated for 16 hr. Cells were collected by centrifugation, resuspended in lysis buffer, and flash frozen in liquid nitrogen.

Cell pellets were resuspended in Lysis buffer (1X PBS supplemented with 250 mM NaCl, 0.1% Tween-20, 1 mM dithiothreitol, and 1 mM phenylmethylsulfonyl fluoride). Cells were disrupted by sonication, and the lysate was cleared by centrifugation. The lysate was applied to 0.5 mL of glutathione agarose (Sigma-Aldrich) per liter of culture for 1 hr at 4°C. Agarose was washed three times in Lysis Buffer, and proteins were eluted using Elution Buffer (50 mM Tris-HCl, 75 mM KCl, 10 mM reduced glutathione, pH 8.0). For certain proteins, the glutathione S-transferase (GST) purification tag was removed by incubation overnight at 4°C with 1 mg HRV-3C protease (Cheeseman lab) per 50 mL elution fraction. Final polishing was performed by gel-filtration on a Superdex 200 16/60 coupled to an AktaPurifier system (GE Healthcare) into Binding Buffer (50 mM HEPES, 400 mM KCl, 10% glycerol, 0.1% Tween-20, 1 mM EDTA, 1 mM dithiothreitol, pH 7.4). Peak fractions were concentrated in Vivaspin concentrators (GE Healthcare), aliquoted, and snap-frozen in liquid nitrogen.

#### In vitro binding assays and gel filtration

50 μL binding reactions were prepared with recombinant proteins diluted to 3.5 μM each in Binding Buffer (50 mM HEPES, 400 mM KCl, 10% glycerol, 0.1% Tween-20, 1 mM EDTA, 1 mM dithiothreitol, pH 7.4). Reactions were incubated on ice for 1 hr before clearing by centrifugation at >10,000 g for 10 min. Cleared samples were run over a Superdex 200 3.2/300 column on an Akta Micro FPLC system (GE Healthcare) pre-equilibrated in Binding Buffer. 40 μL fractions were analyzed by SDS-PAGE followed by staining with Acquastain (Bulldog Bio). For examination of the binding interaction under variable salt concentrations, the KCl concentration in the Binding Buffer was altered according to Fig. legends.

#### Phospho-antibody generation

The CENP-C pS311 phosphospecific antibody was generated against a synthesized phosphopeptide with the following amino acid sequence: CNLRNEE(pS)VLLFTQ (New England Peptide; Covance). Peptide was coupled to Sulfolink Coupling Resin (Thermo Fisher Scientific). Serum from immunized rabbit was depleted against the unphosphorylated peptide and affinity purified against the phosphorylated peptide.

#### Mouse oocyte injection

Germinal vesicle intact oocytes were microinjected with ~5 pL of cRNAs in M2 medium containing milrinone at room temperature with a micromanipulator TransferMan NK 2 (Eppendorf) and picoinjector (Medical Systems Corp.). Injected oocytes were kept in CZB with milrinone for 6-12 hr to allow protein expression, before switching to milrinone-free CZB medium to develop to meiosis I (6.5 hr in vitro maturation) or meiosis II (16 hr in vitro maturation). Oocytes were checked for germinal vesicle breakdown 1.5 hr after milrinone washout, and those that did not breakdown the germinal vesicle were discarded. Meiosis II eggs were activated to progress to anaphase II in CZB medium containing 5mM SrCl_2_ and 2 mM EGTA for 1 hr as previously described (Kishigami and Wakayama, 2007).

Plasmids for cRNA generation were generated in the pCS2+ backbone (von Dassow et al., 2009) and are described in Table S4. cRNAs were synthesized using the T7 mScript Standard mRNA Production System (Cell Script) and injected at ~125 ng/μL final concentration.

#### Oocyte immunocytochemistry and imaging

For immunocytochemistry, samples were fixed in 2% paraformaldehyde in PBS for 30 min at room temperature. The cells were then permeabilized for 15 min in PBS containing 0.2% Triton X-100, blocked in PBS containing 0.2% immunoglobulin G-free bovine serum albumin and 0.01% Tween-20 for 30 min (blocking solution) and then incubated with the primary antibody for 1 hr at room temperature. After four 15 min washings in blocking solution, samples were incubated for 1 hr with either Alexa Flour 594-conjugated (1:500, Invitrogen) or Cy5-conjugated (1:100, Jackson ImmunoResearch) secondary antibody diluted in blocking solution. After an additional three 15-min washings in blocking solution, the samples were mounted in Vectashield mounting solution containing DAPI (Vector Laboratories). Confocal images were collected with a microscope (DMI4000 B; Leica) equipped with a 63× 1.3 NA glycerol-immersion objective lens, an x-y piezo Z stage (Applied Scientific Instrumentation), a spinning disk confocal scanner (Yokogawa Corporation of America), an electron multiplier charge-coupled device camera (ImageEM C9100-13; Hamamatsu Photonics), and an LMM5 laser merge module with 488- and 593-nm diode lasers (Spectral Applied Research) controlled by MetaMorph software (Molecular Devices). Confocal images were collected as z-stacks at 1 μm intervals to visualize the entire meiotic spindle.

To quantify Plk1 and Bub1 signal intensity at kinetochores, CENP-C staining was used to select kinetochores. For each kinetochore, a maximum intensity projection of the optical Z-sections containing CENP-C signal was performed using Fiji/Image J. A circle was drawn around the CENP-C signal, and the same circle was used to quantify Plk1 or Bub1 signal intensity. The mean signal intensity was measured for each circle after subtracting background signal from the surrounding area.

### Quantification and statistical analysis

Fiji/ImageJ (NIH) was used for image manipulation and kinetochore quantification. Where indicated in the method details, a custom CellProfiler pipeline was used. Statistical tests and analysis of enzyme kinetics were performed in Graphpad Prism. All statistical details including type of test and exact value of n is included in the figure legends.

---

**Table S1.**
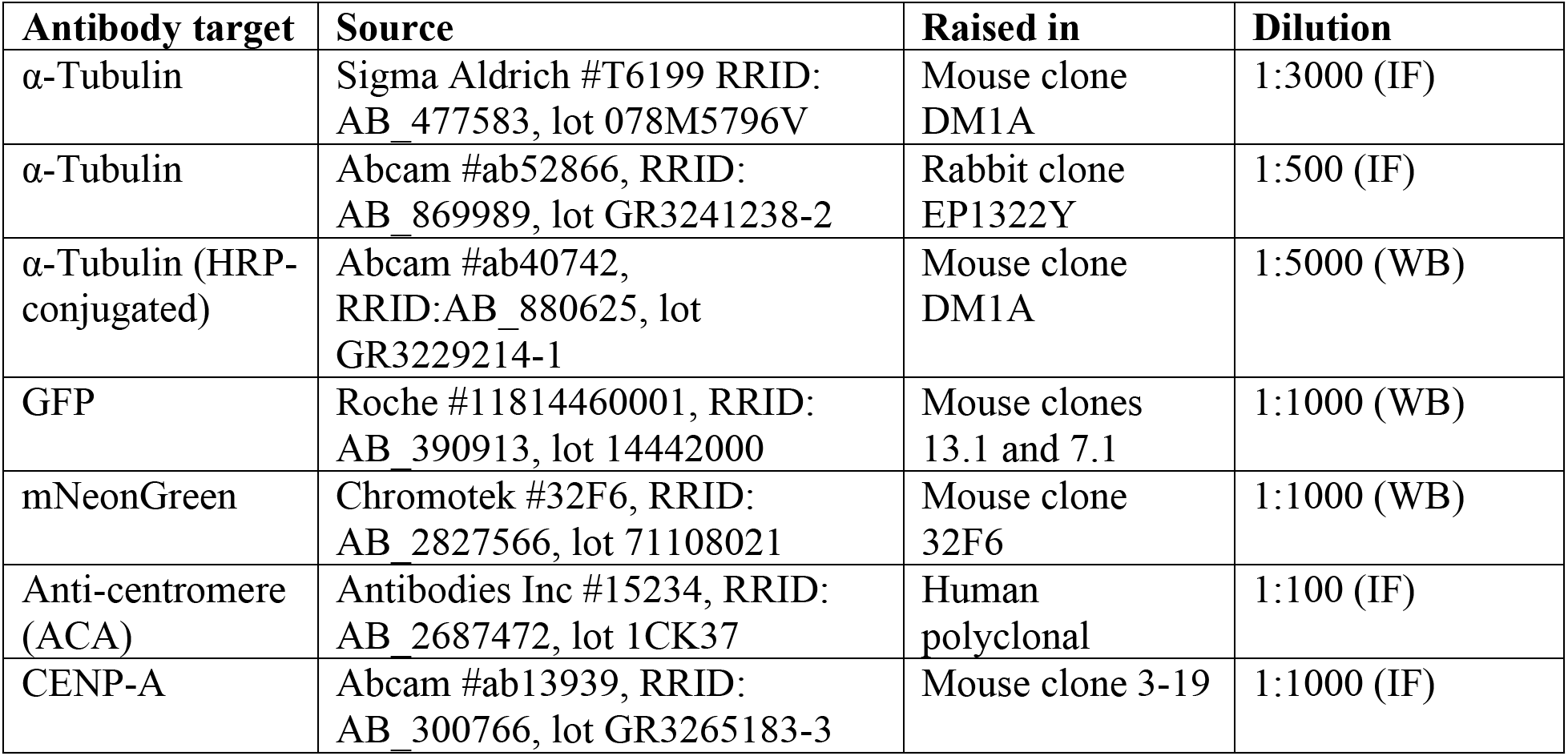
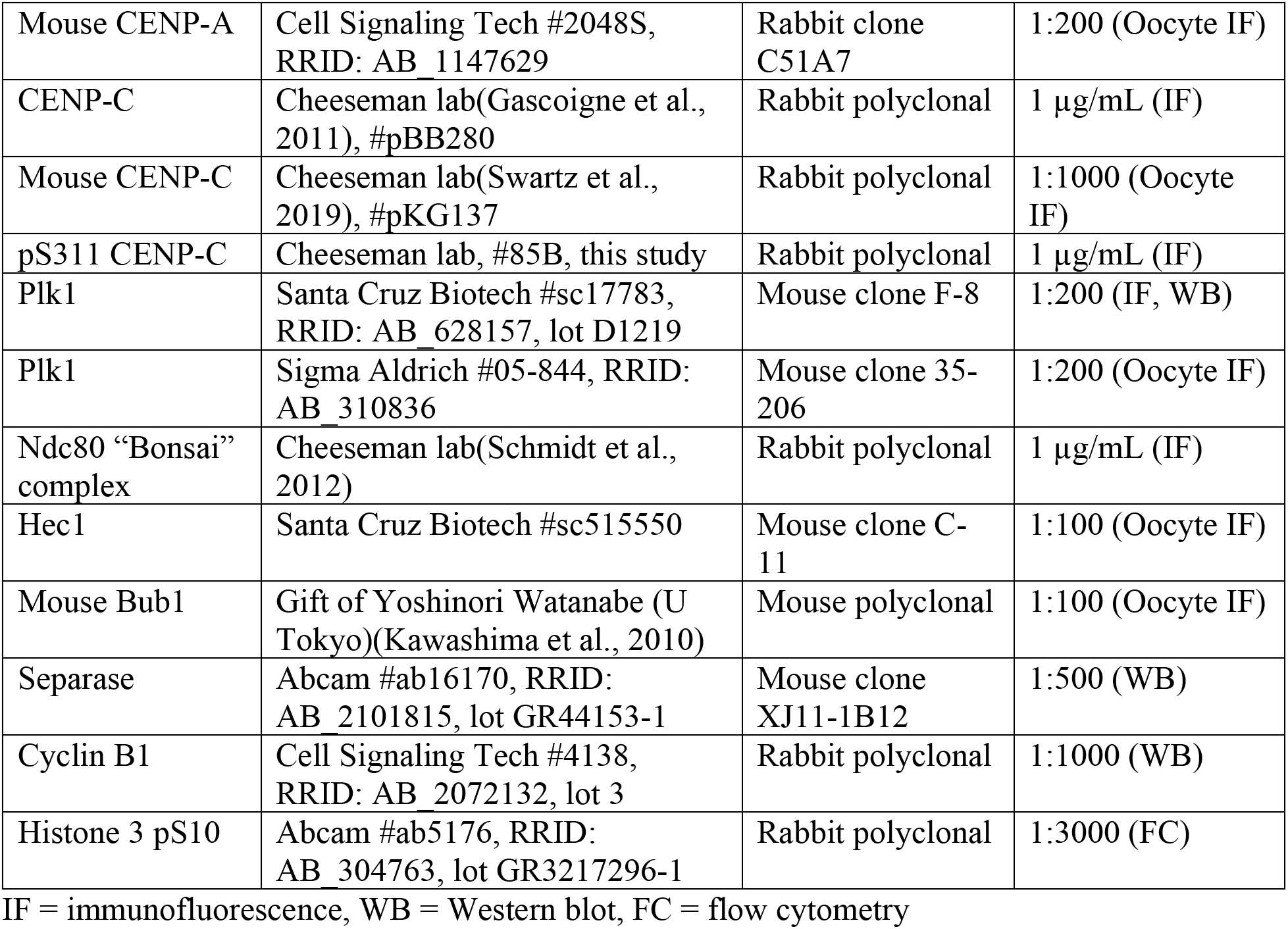
Antibodies used in this study.

**Table S2.** Cell lines used in this study.

| **Name** | **Description** | **Expression** | **Source** |
| --- | --- | --- | --- |
| cNM216-7 | 2xmNeonGreen-hMeikin | Constitutive | Retroviral transduction with pNM481, clonal |
| cNM239-14 | 2xmNeonGreen-hMeikin (328-373) | Constitutive | Retroviral transduction with pNM512, clonal |
| cNM240-4 | 2xmNeonGreen-hMeikin (1-332) | Constitutive | Retroviral transduction with pNM513, clonal |
| cNM238-3 | 2xmNeonGreen-hMeikin (ICCII/DIII mutant) | Constitutive | Retroviral transduction with pNM511, clonal |
| cNM396-15 | hMeikin-mNeonGreen | Constitutive | Retroviral transduction with pNM795, clonal |
| cNM087 | EGFP | Dox-Inducible | AAVS1 Safe-harbor insertion of pNM280 |
| cNM102 | EGFP-hMeikin | Dox-Inducible | AAVS1 Safe-harbor insertion of pNM299 |
| cNM220 | EGFP-hMeikin (104-373) | Dox-Inducible | AAVS1 Safe-harbor insertion of pNM480 |
| cNM244 | EGFP-hMeikin (124-373) | Dox-Inducible | AAVS1 Safe-harbor insertion of pNM549 |
| cNM271 | EGFP-hMeikin (155-373) | Dox-Inducible | AAVS1 Safe-harbor insertion of pNM603 |
| cNM142 | EGFP-hMeikin (184-373) | Dox-Inducible | AAVS1 Safe-harbor insertion of pNM335 |
| cNM143 | EGFP-hMeikin (199-373) | Dox-Inducible | AAVS1 Safe-harbor insertion of pNM336 |
| cNM134 | EGFP-hMeikin (227-373) | Dox-Inducible | AAVS1 Safe-harbor insertion of pNM325 |
| cNM123 | EGFP-hMeikin (1-332) | Dox-Inducible | AAVS1 Safe-harbor insertion of pNM310 |
| cNM145 | EGFP-hMeikin (1-310) | Dox-Inducible | AAVS1 Safe-harbor insertion of pNM338 |
| cNM124 | EGFP-hMeikin (1-301) | Dox-Inducible | AAVS1 Safe-harbor insertion of pNM311 |
| cNM226 | EGFP-hMeikin (1-226) | Dox-Inducible | AAVS1 Safe-harbor insertion of pNM502 |
| cNM311 | EGFP-hMeikin (1-154) | Dox-Inducible | AAVS1 Safe-harbor insertion of pNM680 |
| cNM273 | EGFP-hMeikin (124-332) | Dox-Inducible | AAVS1 Safe-harbor insertion of pNM611 |
| cNM172 | EGFP-hMeikin (175-181 4A mut) | Dox-Inducible | AAVS1 Safe-harbor insertion of pNM400 |
| cNM170 | EGFP-hMeikin (S196A) | Dox-Inducible | AAVS1 Safe-harbor insertion of pNM398 |
| cNM119 | EGFP-hMeikin (STP mutant) | Dox-Inducible | AAVS1 Safe-harbor insertion of pNM300 |
| cNM272 | EGFP-hMeikin (8A mutant) | Dox-Inducible | AAVS1 Safe-harbor insertion of pNM604 |
| cNM321 | EGFP-hMeikin (E151R, R154E) | Dox-Inducible | AAVS1 Safe-harbor insertion of pNM726 |
| cNM371 | EGFP-hMeikin (1-332)-H2B | Dox-Inducible | AAVS1 Safe-harbor insertion of pNM801 |
| cNM469 | EGFP-hMeikin (1-154)-H2B | Dox-Inducible | AAVS1 Safe-harbor insertion of pNM888 |
| cNM366 | EGFP-hMeikin (155-332)-H2B | Dox-Inducible | AAVS1 Safe-harbor insertion of pNM802 |
| cNM370 | EGFP-hMeikin (1-332, STP mutant)-H2B | Dox-Inducible | AAVS1 Safe-harbor insertion of pNM804 |
| cNM372 | EGFP-hMeikin (1-332, 8A mut)-H2B | Dox-Inducible | AAVS1 Safe-harbor insertion of pNM803 |
| cNM426e | H2B-mScarlet-hMeikin(1-332)-mNeonGreen | Constitutive | Retroviral transduction with pNM848, fluorescence enriched |
| cNM427e | H2B-mScarlet-hMeikin(1-332, E151R, R154E)-mNeonGreen | Constitutive | Retroviral transduction with pNM849, fluorescence enriched |
| cNM428e | H2B-mScarlet-hMeikin(1-332, S149A)-mNeonGreen | Constitutive | Retroviral transduction with pNM850, fluorescence enriched |
| cNM430e | H2B-mScarlet-hMeikin(1-332, 8A mut)-mNeonGreen | Constitutive | Retroviral transduction with pNM852, fluorescence enriched |
| cNM431e | H2B-mScarlet-hRad21(142-476)-mNeonGreen | Constitutive | Retroviral transduction with pNM853, fluorescence enriched |
| cNM432e | H2B-mScarlet-hRad21(142-476, E169R, R172E, E447R, R450E, E457R, R460E)-mNeonGreen | Constitutive | Retroviral transduction with pNM854, fluorescence enriched |
| cNM478e | H2B-mScarlet-hRec8(297-506)-mNeonGreen | Constitutive | Retroviral transduction with pNM937, fluorescence enriched |
| cNM486e | H2B-mScarlet-hRec8(297-506)-hMeikin(1-332, E151R, R154E)-mNeonGreen | Constitutive | Retroviral transduction with pNM869, fluorescence enriched |
| cNM500e | H2B-mScarlet-hRec8(297-506, E401R, R404E)-hMeikin(1-332, E151R, R154E)-mNeonGreen | Constitutive | Retroviral transduction with pNM972, fluorescence enriched |
| cNM487e | H2B-mScarlet-hRec8(297-506)-hMeikin(155-332)-mNeonGreen | Constitutive | Retroviral transduction with pNM870, fluorescence enriched |
| cNM493e | H2B-mScarlet-hRec8(297-506)-hMeikin(1-332, E151R, R154E, 8A mut)-mNeonGreen | Constitutive | Retroviral transduction with pNM946, fluorescence enriched |

All cell lines listed are derived from HeLa cells

**Table S3.** Recombinant Proteins used in this study.

| **Protein** | **Plasmid** |
| --- | --- |
| GST-CENP-C (700-943) | pKM138 |
| GST-CENP-C (808-943) | pNM221 |
| GST-CENP-C (775-943, 835-838A) | pNM232 |
| GST-CENP-C (775-943, 865-867A) | pNM271 |
| *GST-sfGFP-Meikin (328-373) | pNM190 |
| *GST-sfGFP-Meikin (328-373, ICCII/DIII mutant) | pNM201 |
| *GST-sfGFP-Meikin (302-373) | pNM175 |

\* = GST-tag removed during purification

**Table S4.**
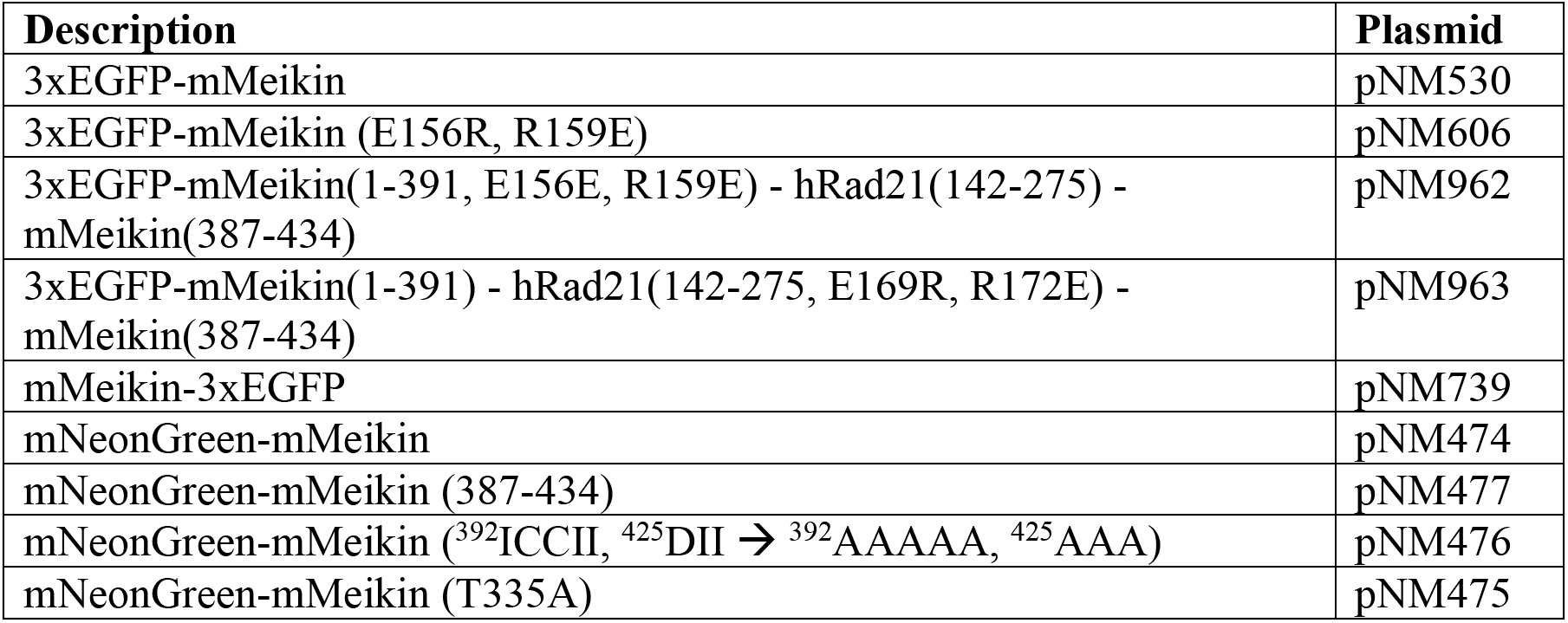
Plasmids used for oocyte experiments.

## Supplemental Figure Legends

**Supplemental Figure 1.**
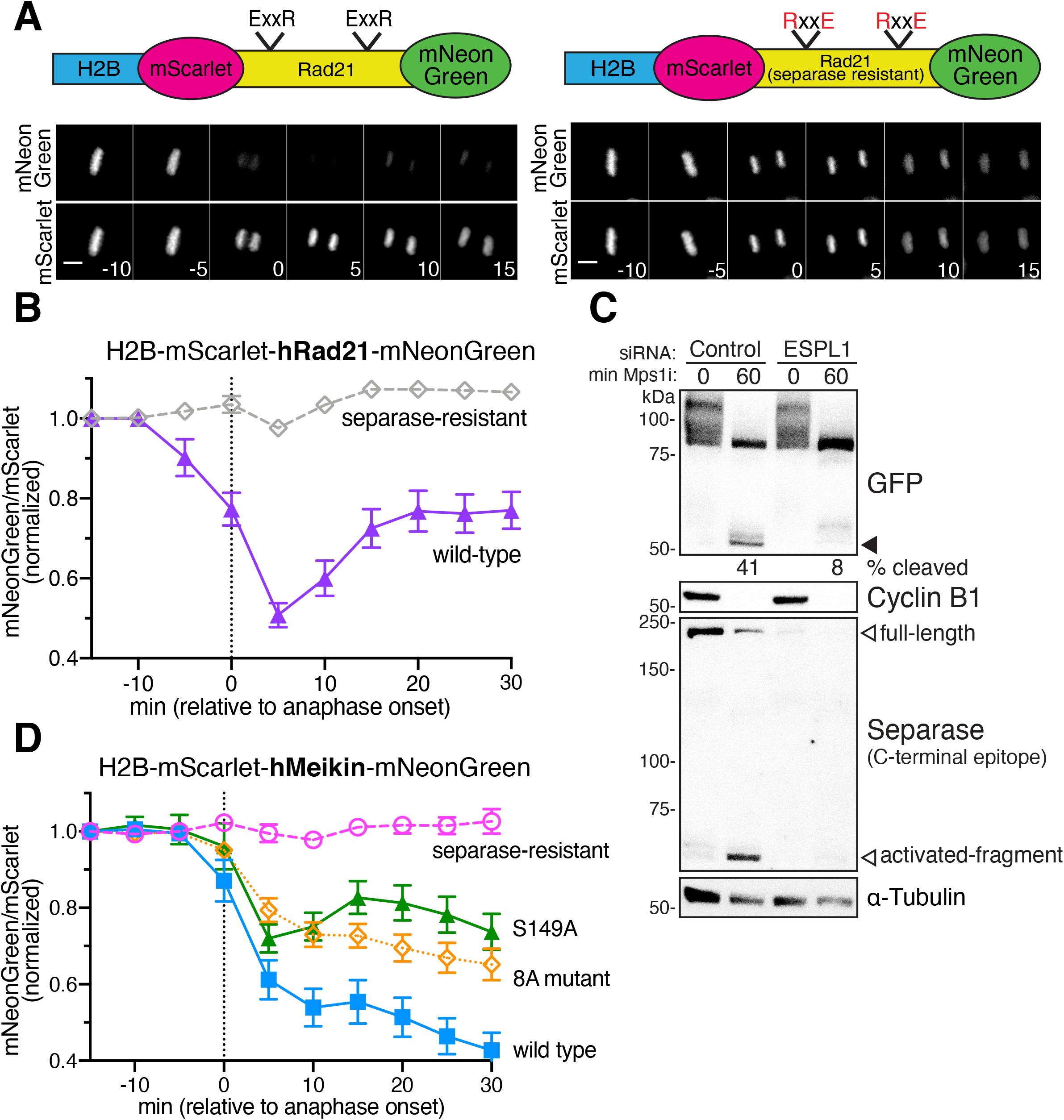
(Related to Figure 1): Dual color H2B-cleavage sensors monitor Separase activity at anaphase. **A.** Schematic of dual-color hRad21 H2B-cleavage sensor. A fragment of hRad21 (amino acids 142-476) including the predicted Separase cleavage sites was used. Proteolytic cleavage of Rad21 leads to the release and diffusion of C-terminal mNeonGreen, but retention of the N-terminal H2B-mScarlet on the DNA. Montage of time lapse images of cells expressing the indicated sensor. Numbers indicate minutes relative to anaphase onset. **B.** Quantification of the mNeonGreen:mScarlet ratio of Rad21 H2B cleavage sensors. The increase in mNeonGreen signal after +10 min in the wild-type sensor coincides with DNA decondensation and likely reformation of the nuclear envelope. Thus, we hypothesize that this increase is due to nuclear import of the cleaved C-terminal fragment. **C.** HeLa cells were treated with the indicated siRNA for 24 hr then induced to express EGFP-hMeikin by treatment with doxycycline and arrested in mitosis by treatment with nocodazole. Cells were forced to exit from mitosis by treatment with Mps1i. At the indicated timepoint, cells were collected, lysed, and analyzed by Western blotting. Percent cleavage was calculated by densitometry analysis of the GFP blot. **D.** Quantification of the mNeonGreen:mScarlet ratio of Meikin mutant H2B cleavage sensors as in Fig. 1E. The number of cells analyzed per condition were: Rad21 (19 cells), Rad21(separase-resistant) (23 cells), Meikin(S149A) (15 cells), Meikin(8A mutant) (23 cells). Data for wild type (15 cells) and Separase-resistant (15 cells) Meikin sensors are duplicated from Fig. 1. Error bars indicate 95% confidence intervals. Scale bars, 10 μm.

**Supplemental Figure 2.**
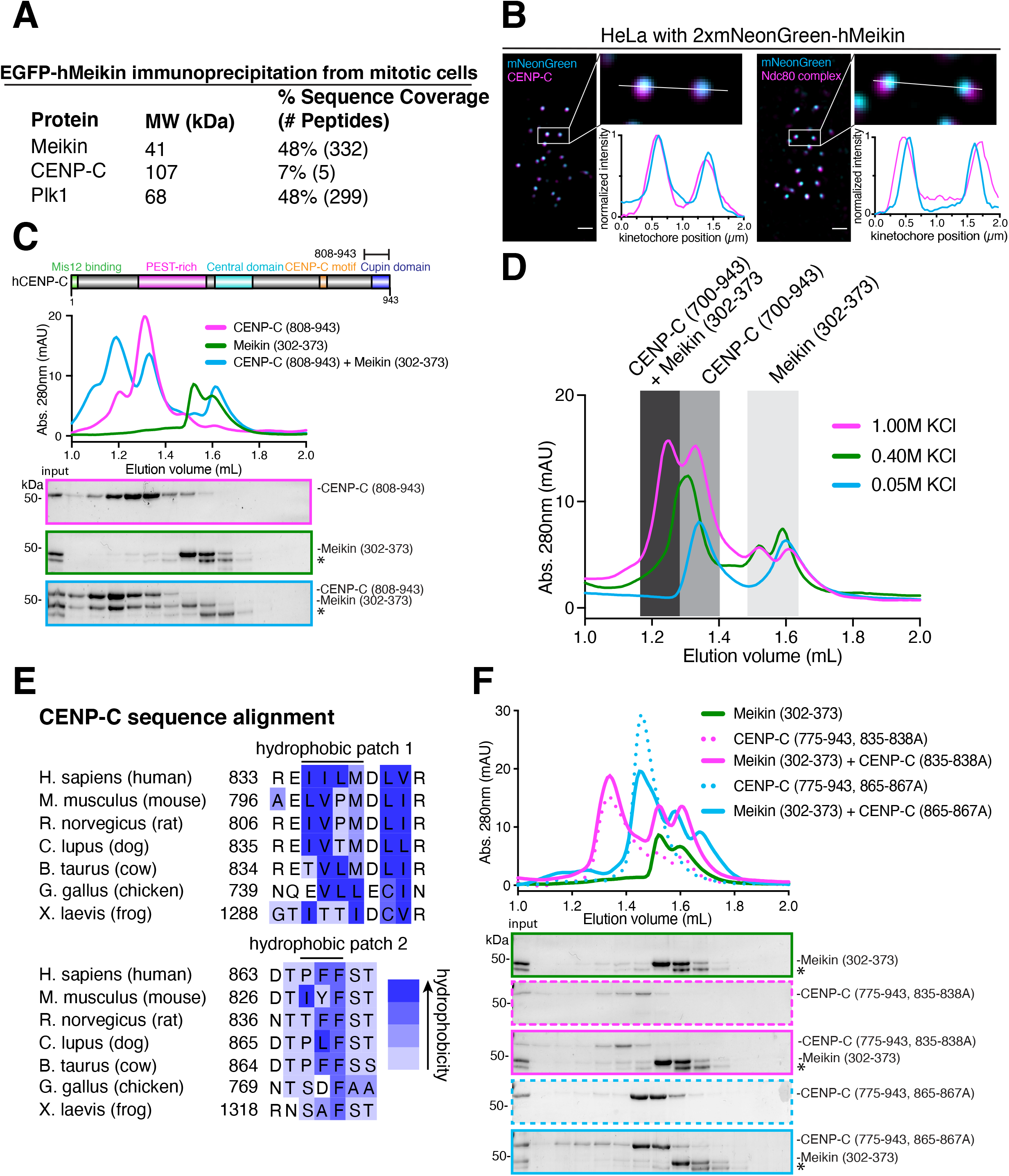
(Related to Figure 2): Meikin binds to CENP-C through hydrophobic patches on both proteins. **A.** GFP-immunoprecipitates from HeLa cells expressing EGFP-hMeikin were analyzed by mass-spectrometry. The percent sequence coverage and number of total peptides of the indicated proteins is shown. Data is the sum of multiple mass-spectrometry experiments. **B.** Metaphase HeLa cells stably expressing 2xmNeonGreen-hMeikin were fixed in ice cold methanol for 10 min then stained for CENP-C or the Ndc80 complex. Linescan analysis of the boxed sister-kinetochore pairs is shown. Meikin localizes to the inner kinetochore overlapping with CENP-C. Scale bar is 1 μm. Deconvolved immunofluorescence images are shown. Linescan analysis was performed before deconvolution. **C.** Diagram of known kinetochore protein interaction sites within CENP-C. Recombinant sfGFP-tagged Meikin and GST-tagged CENP-C protein fragments were bound and complexes analyzed by gel filtration. Fractions corresponding to elution volumes of 1.0 to 2.0 mL were analyzed by SDS-Page and Coomassie staining. **D.** Recombinant sfGFP-tagged Meikin and GST-tagged CENP-C protein fragments were bound and complexes analyzed by gel filtration. Binding reactions and gel filtration were performed in buffer with the indicated KCl concentration. **E.** Sequence alignment of CENP-C from selected vertebrates with conserved hydrophobic patches indicated. Amino acid hydrophobicity is indicated in blue. **F.** Recombinant Meikin and CENP-C fragments containing mutations in conserved hydrophobic patches analyzed by gel filtration as above.

**Supplemental Figure 3.**
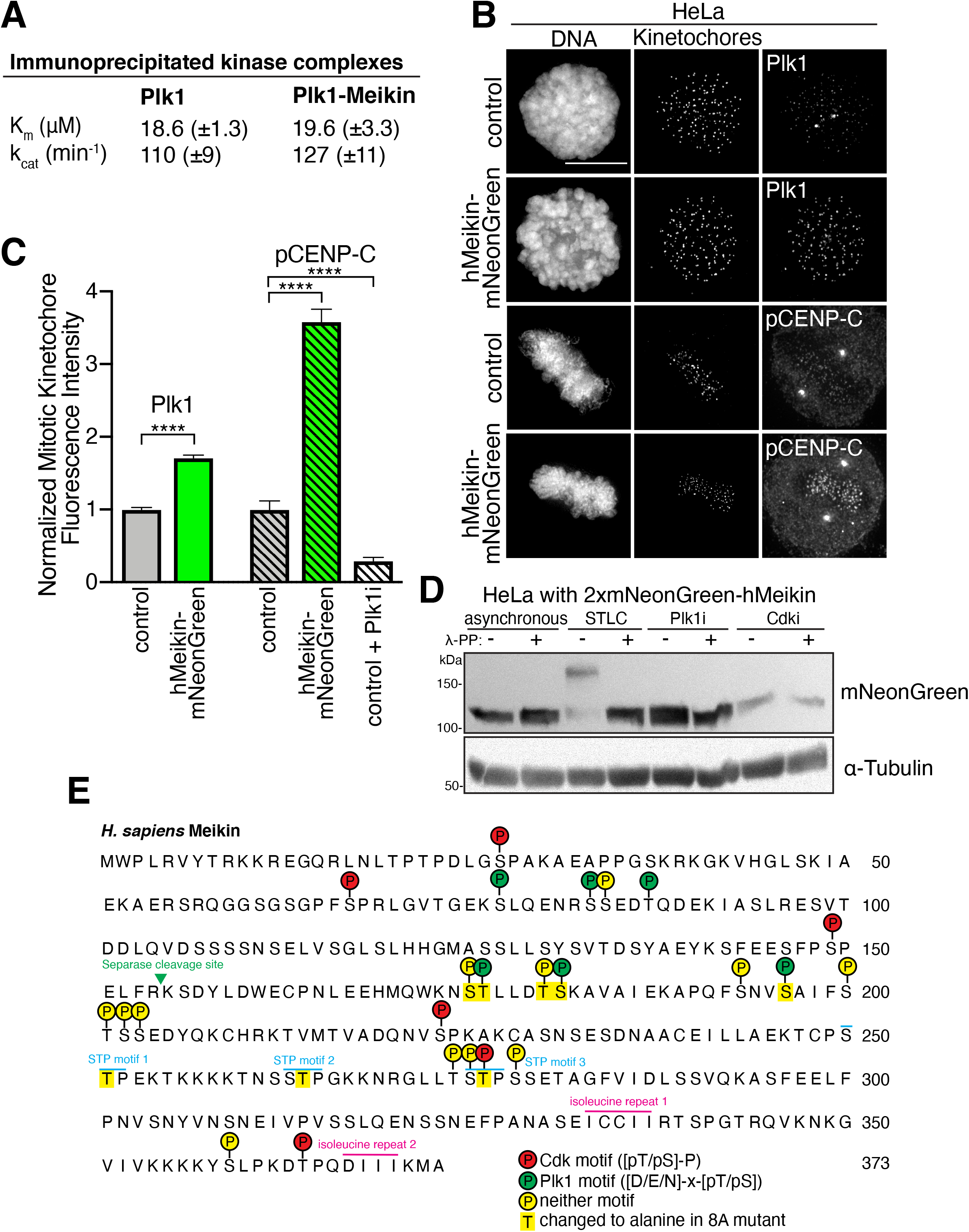
(Related to Figure 3): Meikin binds active Plk1. **A.** Michaelis-Menton parameters derived from kinase assays of Plk1 or Plk1-Meikin complexes immunoprecipitated from HeLa cells. Standard error is represented in brackets. **B.** Deconvolved immunofluorescence images of HeLa cells (control) or HeLa cells stably expressing hMeikin-mNeonGreen and stained for Plk1 (pre-extracted in PBS + 0.5% Triton-X100 for 5 min; pre-extraction disrupts the bipolar spindle and centrosome staining of Plk1; kinetochores are co-stained with CENP-C antibody) or phosphorylated S311 on CENP-C (fixed in ice-cold methanol at −20°C for 20 min; centrosome pCENP-C staining is non-specific; kinetochores are co-stained with CENP-A antibody). Images of similarly stained cells are scaled identically. Scale bars, 10 μm. **C.** Quantification of kinetochore intensity of mitotic HeLa cells stably expressing hMeikin-mNeonGreen. Values were normalized to the mean of the control. Means and 95% confidence intervals are presented. Specificity of the pS311 CENP-C antibody was demonstrated by treatment of cells with Plk1i for 2 hr prior to staining. \*\*\*\**P* < 0.0001, two-tailed *t*-test. n = Plk1: control (27 cells, 2,911 kinetochores), hMeikin (22 cells, 2,626 kinetochores); control vs hMeikin (t = 29.24, df = 5535). n = pCENP-C: control (21 cells, 1,093 kinetochores), hMeikin (27 cells, 1,185 kinetochores), control + Plk1i (25 cells, 2,302 kinetochores); control vs hMeikin (t = 23.55, df = 2276); control vs control + Plk1i (t = 12.50, df = 3393). Quantification was performed before deconvolution. **D.** HeLa cells stably expressing 2xmNeonGreen-hMeikin were treated with the indicated inhibitors. Whole cell lysates were incubated with or without lambda-phosphatase and analyzed by Western blot. **E.** GFP-immunoprecipitates from mitotic HeLa cells expressing EGFP-hMeikin were analyzed by mass-spectrometry. Phosphorylation sites identified across multiple experiments are indicated on the Meikin protein sequence.

**Supplemental Figure 4.**
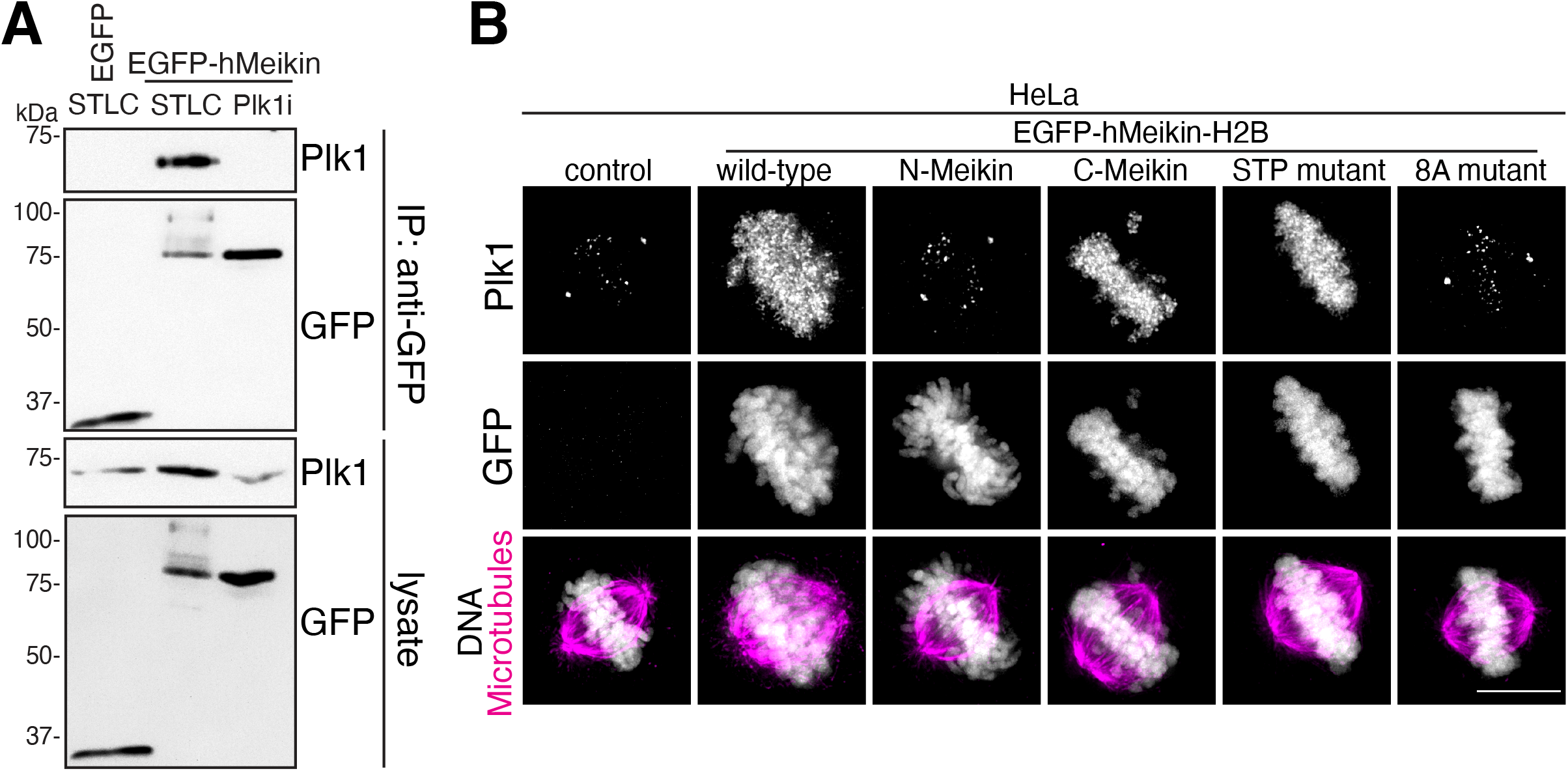
(Related to Figure 3): Full length Meikin and the C-Meikin cleavage fragment bind Plk1. **A.** HeLa cells expressing EGFP-hMeikin were induced with doxycycline and arrested in mitosis by STLC or Plk1i treatment. Cells were lysed and GFP-immunoprecipitation (IP) was performed and analyzed by Western blot. **B.** Deconvolved immunofluorescence images of HeLa cells induced to express EGFP-hMeikin-H2B by treatment with doxycycline. A Meikin fragment excluding the C-terminal kinetochore binding domain was used. Cells were stained for Plk1. Images are not scaled equivalently. Scale bars, 10 μm.

**Supplemental Figure 5.**
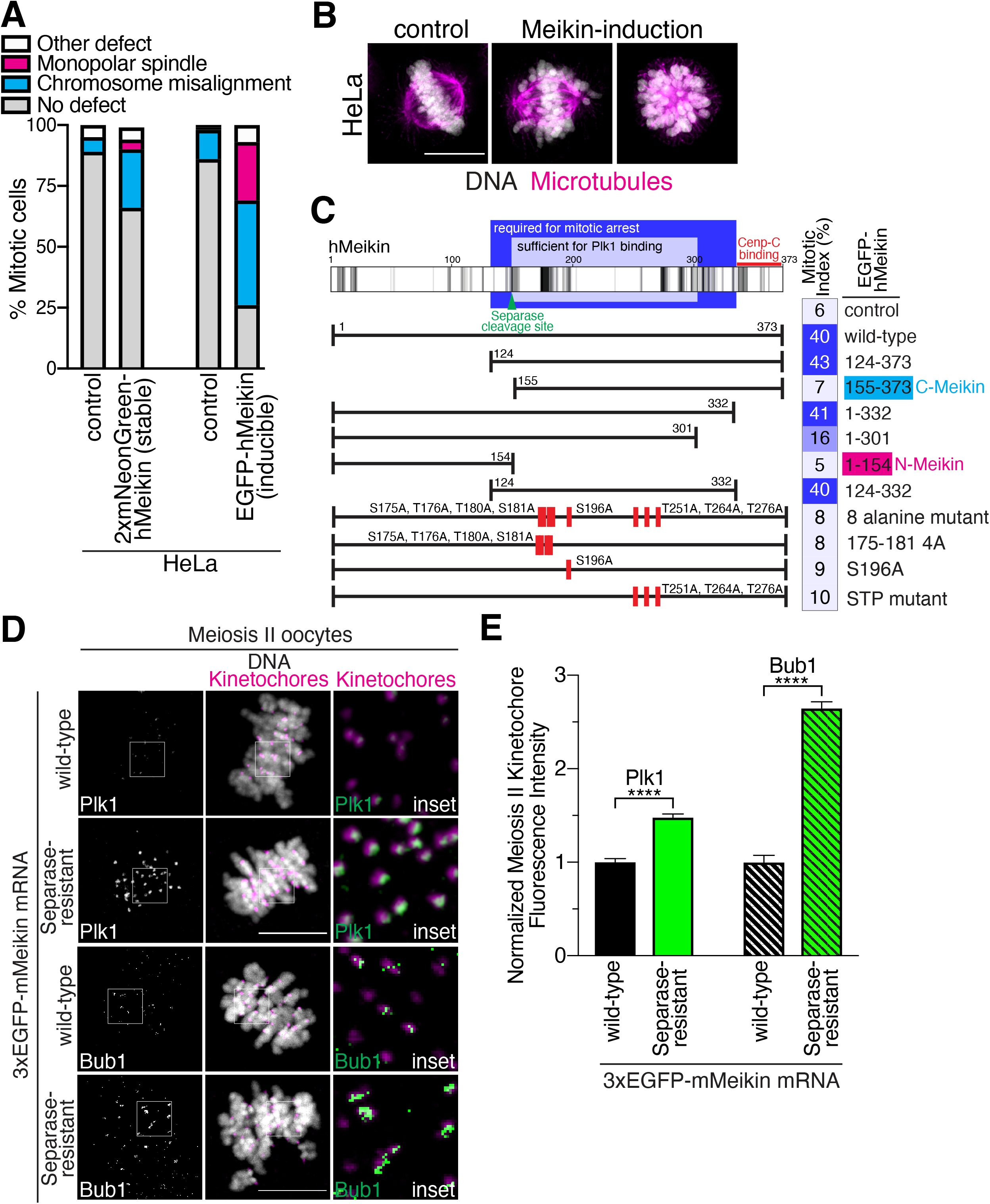
(Related to Figure 4): Meikin expression causes chromosome misalignment and monopolar spindles. **A.** Quantification of mitotic defects observed in HeLa cells stably expressing Meikin or acutely induced to express Meikin from a doxycycline-inducible promoter. 100 mitotic cells were analyzed for each condition. **B.** Representative immunofluorescence images of mitotic HeLa cells stained for DNA and microtubules. High Meikin levels were induced in HeLa cells using a doxycycline inducible promoter. Scale bar, 10 μm. **C.** Schematic of the human Meikin protein with conserved residues (as measured by Consurf(Ashkenazy et al., 2016)) indicated in black. The indicated EGFP-Meikin constructs were expressed in HeLa cells by doxycycline induction. Cells were fixed and stained for the mitotic marker phosphorylated histone 3. The mitotic index of GFP-positive cells was measured by flow cytometry. The minimal region sufficient for Plk1 binding and for the mitotic arrest phenotype are indicated. **D.** Mouse oocytes injected with the indicated 3xEGFP-mMeikin mRNA were matured to meiosis II and stained for Plk1 or Bub1. Kinetochores are stained with mouse CENP-C antibody. Images of similarly stained cells are scaled identically. Insets, 5 μm. Scale bars, 10 μm. **E.** Quantification of kinetochore intensity of Plk1 and Bub1 in meiosis II oocytes injected with the indicated mMeikin mRNA. Values were normalized to the mean of wild-type. Means and 95% confidence intervals are presented. \*\*\*\**P* < 0.0001, two-tailed *t*-test. n = Plk1 (pool of 2 independent experiments, t = 16.53, df = 890): wild-type (362 kinetochores), Separase-resistant (530 kinetochores), Bub1 (pool of 3 independent experiments, t = 29.55, df = 1344): wild-type (449 kinetochores), Separase-resistant (897 kinetochores).

